# Karyotypic divergence confounds cellular phenotypes in large pharmacogenomic studies

**DOI:** 10.1101/574350

**Authors:** Rene Quevedo, Nehme El-Hachem, Petr Smirnov, Zhaleh Safikhani, Trevor J. Pugh, Benjamin Haibe-Kains

## Abstract

**Background:** Somatic copy-number alterations that affect large genomic regions are a major source of genomic diversity in cancer and can impact cellular phenotypes. Clonal heterogeneity within cancer cell lines can affect phenotypic presentation, including drug response.

**Methods:** We aggregated and analyzed SNP and copy number profiles from six pharmacogenomic datasets encompassing 1,691 cell lines screened for 13 molecules. To look for sources of genotype and karyotype discordances, we compared SNP genotypes and segmental copy-ratios across 5 kb genomic bins. To assess the impact of genomic discordances on pharmacogenomic studies, we assessed gene expression and drug sensitivity data for compared discordant and concordant lines.

**Results:** We found 6/1,378 (0.4%) cell lines profiled in two studies to be discordant in both genotypic and karyotypic identity, 51 (3.7%) discordant in genotype, 97 (7.0%) discordant in karyotype, and 125 (9.1%) potential misidentifications. We highlight cell lines REH, NCI-H23 and PSN1 as having drug response discordances that may hinge on divergent copy-number q

**Conclusions:** Our study highlights the low level of misidentification as evidence of effective cell line authentication standards in recent pharmacogenomic studies. However, the proclivity of cell lines to acquire somatic copy-number variants can alter the cellular phenotype, resulting in a biological and predictable effects on drug sensitivity. These findings highlight the need for verification of cell line copy number profiles to inform interpretation of drug sensitivity data in biomedical studies.

## INTRODUCTION

Human cancer cell lines are ubiquitous experimental model for large pharmacogenomics studies seeking to define genomic predictors of drug response ^1–9^. Highly mutable tumors accumulate approximately 8 SNVs every cell division^10^, the majority of which are passenger mutations, while whole chromosome missegregation occur once every 2-5 cell divisions^11–13, 14^ and may affect hundreds to thousands of genes. CNVs occur more frequently ^10–13^ and can affect a larger portion of the genome; ranging from 1000s to >100 million bases per aberration, and can explain as much as 18% of variation in gene expression ^15^. As such, chromosomal instability could result in an unstable cellular phenotype under different conditions and could therefore respond to different drugs.

The International Cell Line Authentication Committee (ICLAC) was formed^16,17^ to set guidelines for cell line usage in biomedical research to mitigate the issue of misidentification, contamination, and instability of genotype and phenotype. While misidentification and contamination are addressed by these standards, heterogeneity and genomic instability remain a problem. As an example, Ben-David et al. reported genetic variability across 106 cell lines with an estimated 19% discordance in non-silent mutations and 26% discordance in CNVs^18^. They also reveal extensive subclonality and heterogeneity within isogenic strains of MCF-7 and A549 impacting cell morphology, doubling time, and drug sensitivity.

Our study explores the genetic instability of cell lines as a means to deconvolute inconsistencies between multiple large pharmacogenomic studies ^6,19–22^. Upon confirming the cellular identities of cancer cell lines used in these studies, we quantified the discordance between molecular profiles and associated these discrepancies with discrepant transcriptomic and pharmacological phenotypes. Overall, we found significant differences in copy-number (CN) profiles within a small panel of cell lines that were associated to differential response to pharmacological compounds in multiple studies. Our results support the notion that the inherent karyotypic instability of cancer cell lines may have an adverse effect on the reproducibility of pharmacological screens.

## METHODS

### Processing of SNP arrays

We analyzed 3,631 Affymetrix Genome-Wide Human SNP 6.0 (Affy6) arrays and 668 HumanOmni2.5-Quad (Omni) arrays from 1,691 cell lines profiled by a compendium of 6 pharmacogenomic studies (Table 1). We downloaded files using accession codes outlined in Table 1 and obtained cell line annotations from PharmacoDB (version 1.0.0) ^23,24^. We removed samples if they failed quality control metrics specific for Affy6 or Omni platforms ^25^. We generated allele-specific copy-number (CN) profiles using HAPSEG/ABSOLUTE^26^ and ASCAT^27^ for Affy6 and Omni datasets respectively (Figure 1). We segmented homologue-specific (HSCR) or total copy ratios (TCR) into 5 kb genomic bins and calculated a variance measure by reprocessing 60% of raw files from each dataset 20 times.

**Figure 1.**
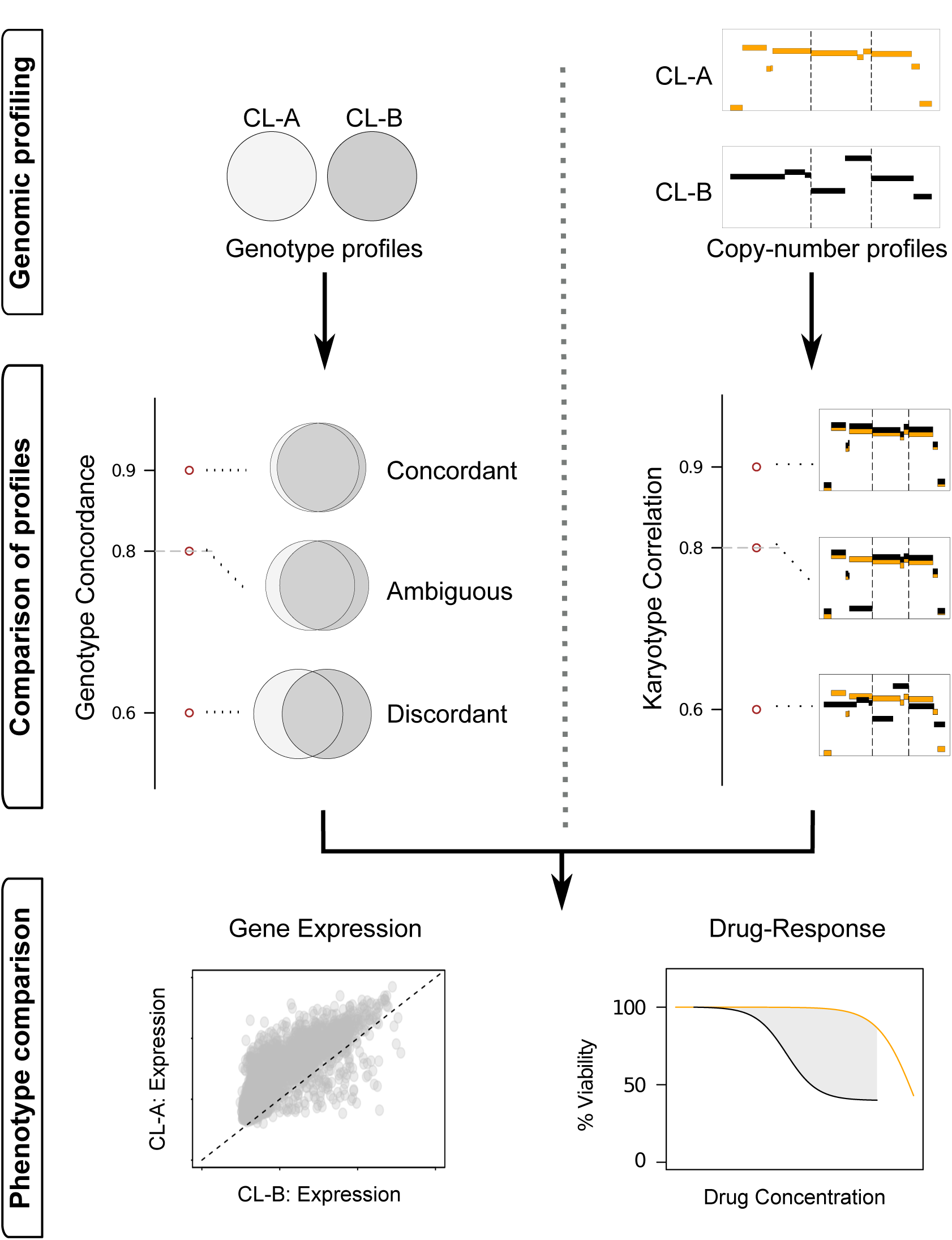
Analytical pipeline used to compare molecular profiles between the GDSC, CGP, CCLE and Pfizer datasets.

**Table 1.**
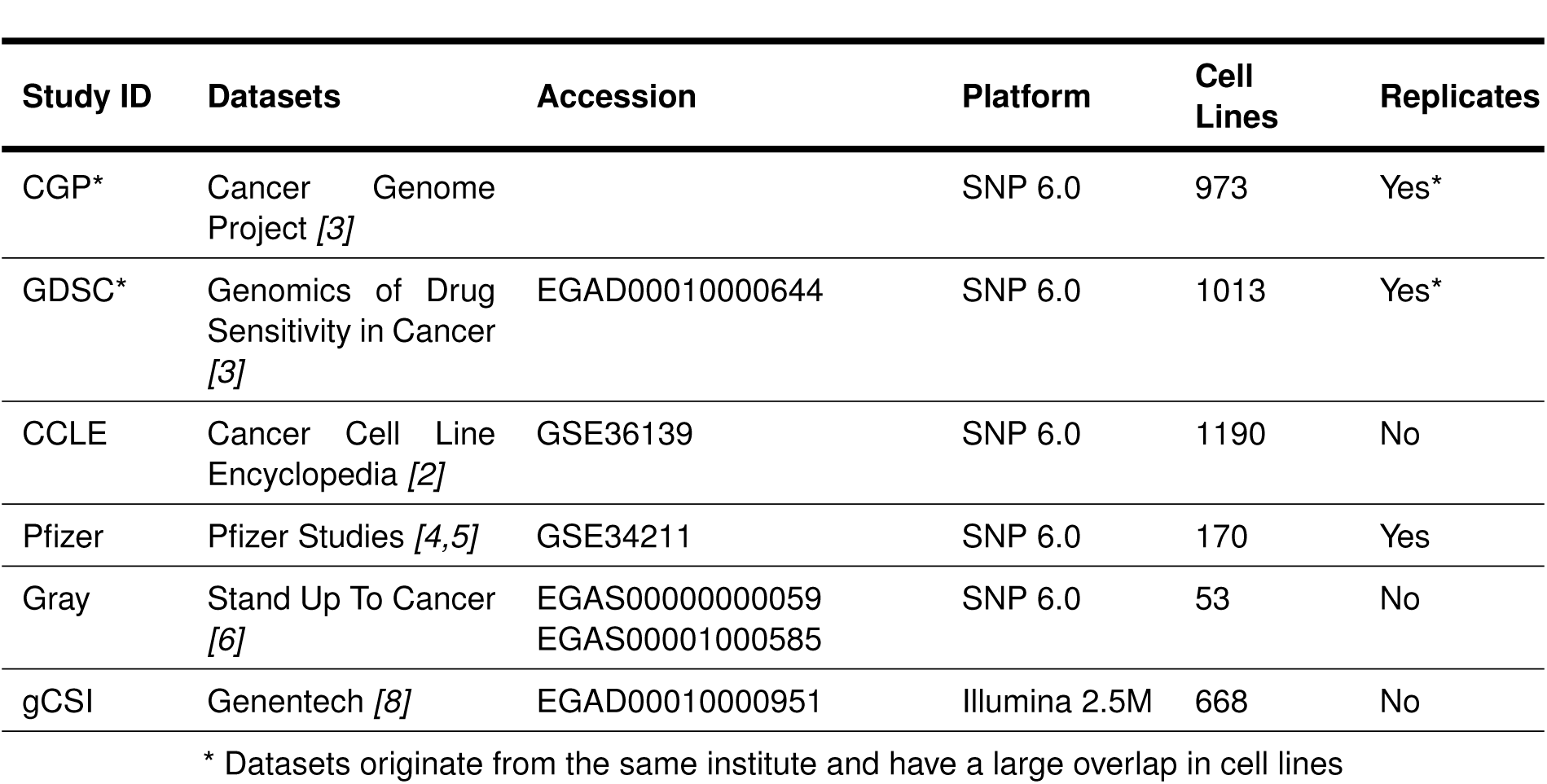
Description of pharmacogenomic datasets

| Study ID | Datasets | Accession | Platform | Cell Lines | Replicates |
| --- | --- | --- | --- | --- | --- |
| CGP* | Cancer Genome Project [3] |  | SNP 6.0 | 973 | Yes* |
| GDSC* | Genomics of Drug Sensitivity in Cancer [3] | EGAD00010000644 | SNP 6.0 | 1013 | Yes* |
| CCLE | Cancer Cell Line Encyclopedia [2] | GSE36139 | SNP 6.0 | 1190 | No |
| Pfizer | Pfizer Studies [4,5] | GSE34211 | SNP 6.0 | 170 | Yes |
| Gray | Stand Up To Cancer [6] | EGAS00000000059<br>EGAS00001000585 | SNP 6.0 | 53 | No |
| gCSI | Genentech [8] | EGAD00010000951 | Illumina 2.5M | 668 | No |

\* Datasets originate from the same institute and have a large overlap in cell lines

### Comparing molecular profiles

To build a reference distribution for concordances between cell line pairs, we built a “matching” distribution consisting of concordance measurements for homonymous cell line pairs. Similarly, we calculated a “non-matching” distribution that consisted of the concordances for heteronymous (i.e. differently named) cell line pairs.

We compared genotypes between two cell lines using a Jaccard index ^28^ (*C*) for all SNPs found on both Affy6 and Omni array platforms. We compared CN-profiles using a variance-weighted Pearson correlation coefficient (r) across all 5 kb genomic bins. We considered genotype and karyotype pairs as matching if the concordance score was greater than 80% or 60% respectively, or if the concordance was within 3 median absolute deviations away from the median (MADM) for the corresponding “matching” cell lines distribution (Figure 1).

We accessed reprocessed drug response curves and gene expression data were accessed from the R PharmacoGx package (version 1.10.3)^23^. Expression space was represented by the landmark genes defined in the L1000 dataset ^29^. We used the recomputed area above the curve (AAC) were used as measurements of drug sensitivity, where larger AACs represented drug-sensitivity ^22^. We computed the distances between drug dose-response curves as the area between the curves (ABC) fitted on overlapping drug concentrations (Figure 1). When comparing phenotypic profiles to the the reference distributions, we used a ratio of likelihoods for *non-match* to *match* in log_2_ space.

### Research reproducibility

Our code and documentation are open-source and publicly available through the CellLineConcordance GitHub repository (github.com/bhklab/CellLineConcordance). A detailed tutorial describing how to run our pipeline and reproduce our analysis results is available the GitHub repository. A virtual machine reproducing the full software environment is available on Code Ocean. SNP data for the GRAY, CCLE and Pfizer datasets are publicly available while the CGP and GDSC data require access permissions (http://www.ebi.ac.uk/ega/dacs/EGAC00001000000).

## RESULTS

### Datasets

To quantify genetic instability of human cancer cell lines used in large-scale pharmacogenomic studies, we reprocessed publicly available Affymetrix SNP arrays from: the Genomics of Drug Sensitivity in Cancer project (GDSC) and its predecessor Cancer Genome Project (CGP), the Cancer Cell Line Encyclopedia (CCLE) project, a compendium of pharmacogenomic projects spearheaded by Pfizer Inc. (Pfizer), and a comprehensive panel of breast cancer cell lines used in the Gray laboratory at Oregon Health & Science University (Gray) (Table 1). We removed the Gray dataset from our analysis as we were unable to reliably differentiate between different zygosity states of SNPs in 28 of the 53 samples (Supp. Fig. 1). An additional 29 of 1,013 samples in GDSC, 66 of 973 in CGP and 5 of 1,190 in CCLE samples were also removed based on the same quality control metrics.

### Cell line authentication

We screened for all isogenic cell line pairs between and within datasets and identified which ones had heteronymous annotations. We described possible relationships using four different classifications: *match* and *possible-match* when cell lines originated or very likely originated from the same patient or sample (e.g., the Burkitt’s lymphoma cell lines EB1 and EB2 from a 7-year old and 9-year old female respectively with no statement explicitly relating the two); *unknown* when no annotations were available to make an informed classification; and *mismatch* when cell lines originated from different patients.

The *match* and *possible-match* categories consisted of cell lines originating from the same tumour (e.g., AU565 and SK-BR-3), same patient but different tissue sites (e.g., head and neck cancer LB771-HNC and peripheral blood LB771-PBL), sub-clonal origin (e.g., SH-SY5Y and the derivative SK-N-SH), or transformed cells (e.g. J82 and Epstein-Barr Virus transformed J82-EBV). Using this metric, we discovered 171 matching lines and 15 possible matching lines across all datasets (Supp. File 1). This analysis did not reveal any unexpected cell line pairs, which can be viewed as a positive control for the concordance analysis.

We cross-referenced cell lines under the *mismatch* category with the registry of misidentified/contaminated cell lines established by ICLAC ^16^ (Fig. 2a). We assumed that overlap of pairs with the registry are indicative of pre-established contaminations or misidentification, while the absence from the registry suggests errors during handling (Supp. Table 1). As validation, all CCLE isogenic heteronymous pairs were identified on the online CCLE portal as being a metastatic pair, identical lines, or derived from the same patient (Supp. Table 2). We also observed GDSC rectifying 34 erroneous cell lines of the 700 homonymous lines from their preceding CGP dataset. Of the 34, SW403 and NB4 were cases of misidentification, 11 failed quality control, and the rest had low genotype concordance with other datasets (Supp. Fig. 2a,b), resulting in the reprocessed samples having significantly better concordance when compared to CCLE (p=0.001; Wilcoxon rank sum test) (Supp. Fig. 2c,d). These results confirm that GDSC and CCLE actively control the identity of their cell lines and update their data accordingly.

**Figure 2.**
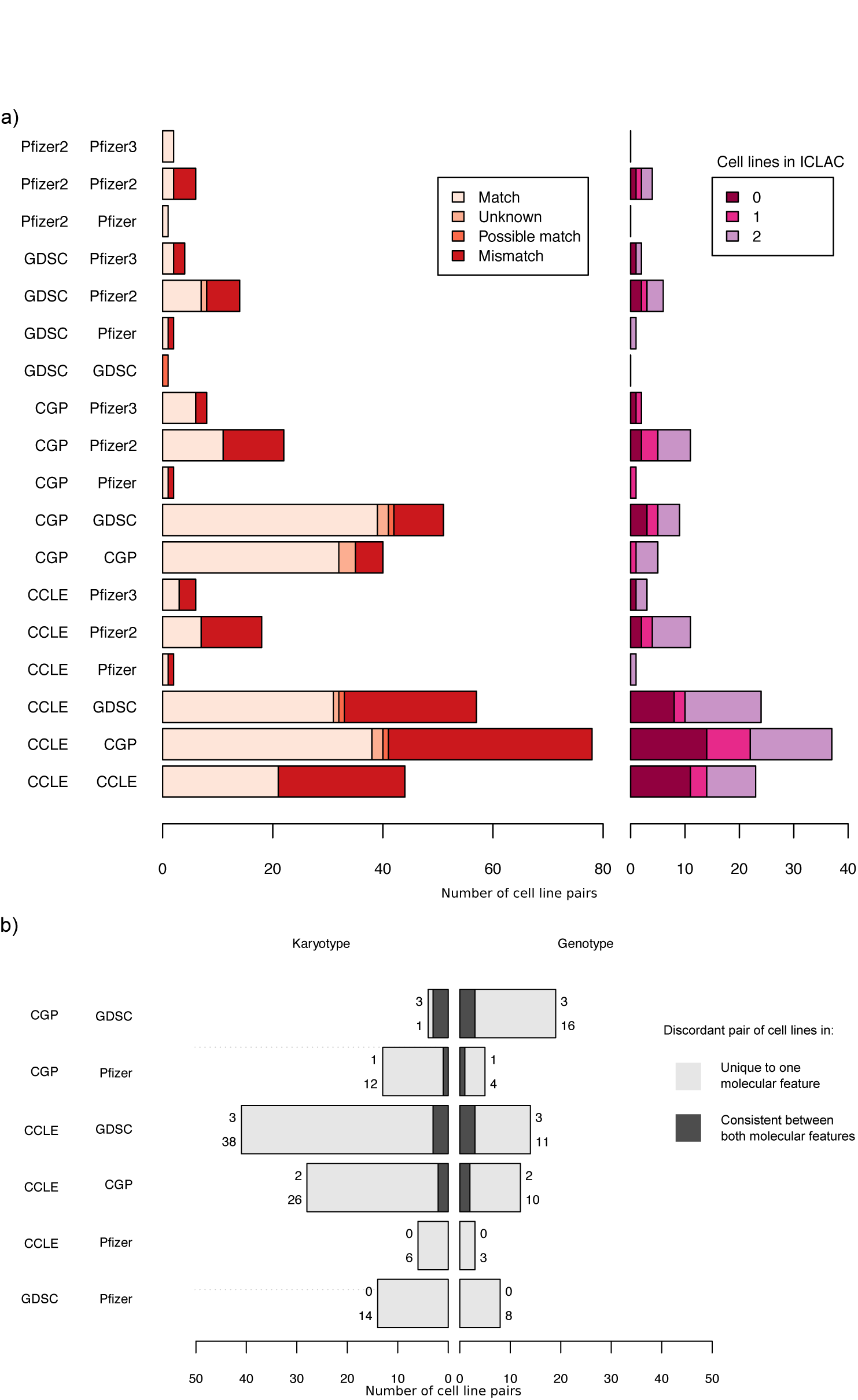
Discrepant cell line pairs with respect to genetic/karyotypic profiles and their annotations. a) Number of cell line pairs that were isogenic despite having different annotations. The cell lines that were isogenic with different annotations were classified as either derived from the same patient (match) or derived from two completely different patients (mismatch). Of the “mismatch” pair of cell lines, we counted whether none (0), one (1), or both (2) of the cell lines were previously reported by ICLAC. b) The number of cell line pairs that were genetically or karyotypically different despite having the same annotations.

### Comparison of karyotypes

We next examined whether isogenic cell lines of the *mismatch* category had concordant karyotypes. The pancreatic ductal adenocarcinoma cell line KCI-MOH1 was flagged as contaminated by HPAC, a cell line of the same cell type. These cell lines represented in CCLE and GDSC respectively were confirmed to be isogenic (concordance *C*=0.902), with concordant karyotypes (r_HSCR-A_: 0.857; r_HSCR-B_: 0.933) and transcriptomes (r_expr_=0.84; Supp. Fig. 3), with a minor shift in log2 ratios on chr2:q24.2-q37.3, chr4, and chr15:q13.2-q26.3 (Fig. 3a). Thus, we concluded that isogenicity of lines is a fair predictor of having similar, but not perfectly concordant molecular profiles.

**Figure 3.**
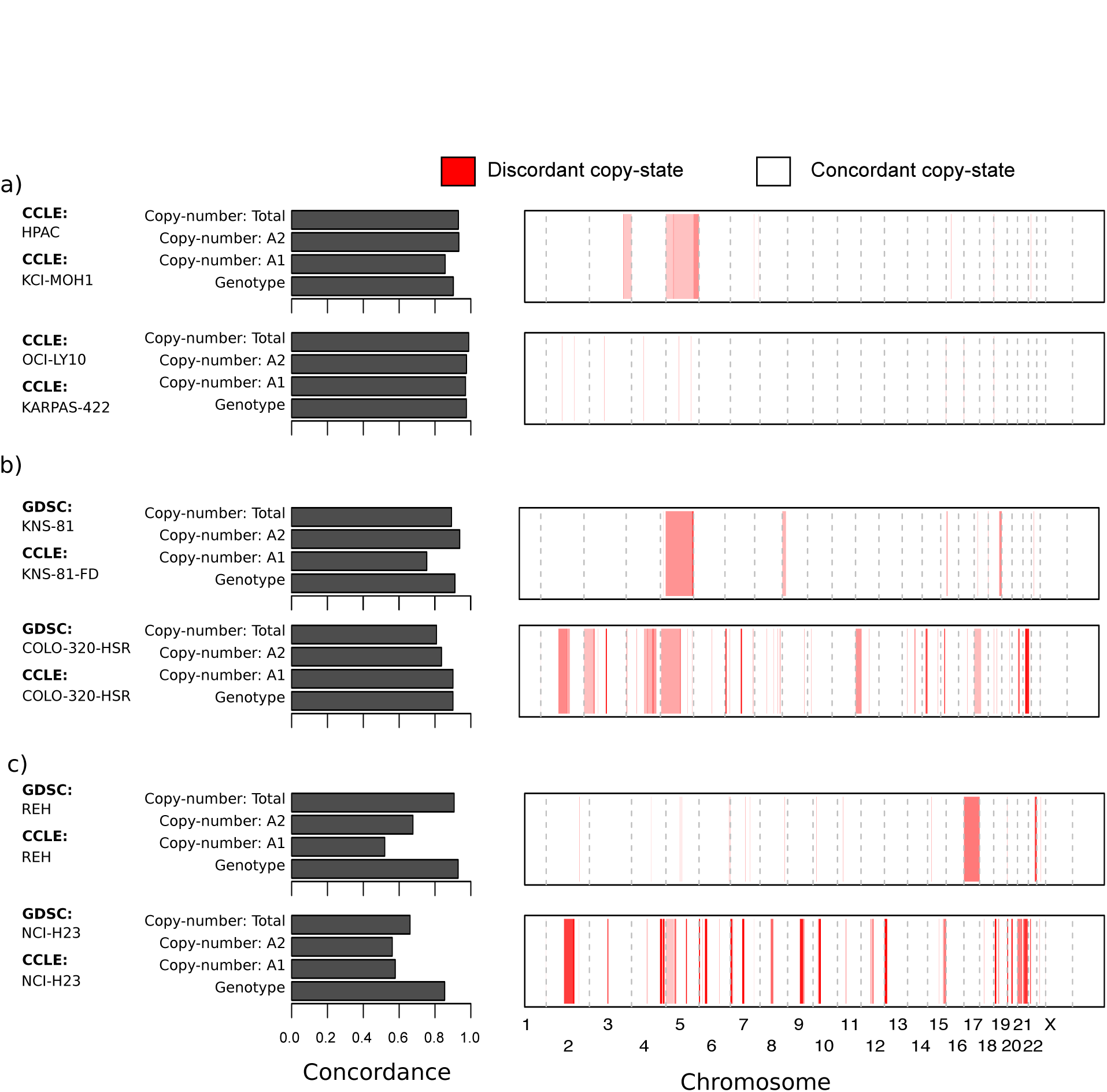
Summarized view of concordance scores and karyotypes between any given two cell lines. Left panel: A summarized view of concordances for genotype and homologue-specific and total copy-number profiles between cell lines. Right panel: Regions of the genome that have a discordant copy-state between any two cell lines. a) Examples of cell lines with discordant annotations but comparable genotypes: HPAC and KCI-MOH1 in CCLE, and OCI-LY10 and KARPAS-422 in CCLE. b) Example of cell lines with expected discordant karyotypes: Transformed cell line KNS-81 and KNS-81-FD, and homologous staining region cell lines COLO-320-HSR. c) Cell lines with unexpected copy-number drift: Low chromosomal instability cell line REH and high chromosomal instability cell line NCI-H23.

Similarly, the two B-cell lymphoma cell lines OCI-LY10 and KARPAS-422 both screened in CCLE were isogenic lines (*C*=0.98) with no history of contamination in ICLAC and presented with comparable karyotypes (r_HSCR-A_=0.97, r_HSCR-B_=0.98; Fig. 3a) and transcriptomes (r_expr_=0.86; Supp. Fig. 3). A comparison of the CCLE screened OCI-LY10 with KARPAS-422 screened in GDSC also displayed comparable molecular profiles (*C*=0.92; r_HSCR-A_: 0.94, r_HSCR-B_: 0.88; and r_expr_=0.86; Supp. Fig. 3). While these two cell lines are of the same cell type, they are derived from two different patients, a 44-year old man and 73-year old female respectively. Since the OCI-LY10 cell line used in CCLE dataset had molecular profiles comparable to KARPAS-422 in both in GDSC and CCLE, we were able to conclude that the OCI-LY10 cell line used in CCLE is likely to be misidentified.

Overall, we observed that the majority of identically annotated cell lines pairs between datasets have either a discordant karyotype or genotype, but not both (Fig. 2b). Cell lines that were discordant in both karyotype and genotype space were largely composed of CGP cell lines that were replaced in the GDSC dataset (Supp. Table 3). While genotype concordance is a binary classification schema for cell line authentication, the karyotype concordance score can be used as a surrogate metric to quantify the amount of CN discrepancies. While correlations greater than 0.8 presented with few to no CN discrepancies, correlations less than 0.6 often had major differences between profiles (Fig. 4).

**Figure 4.**
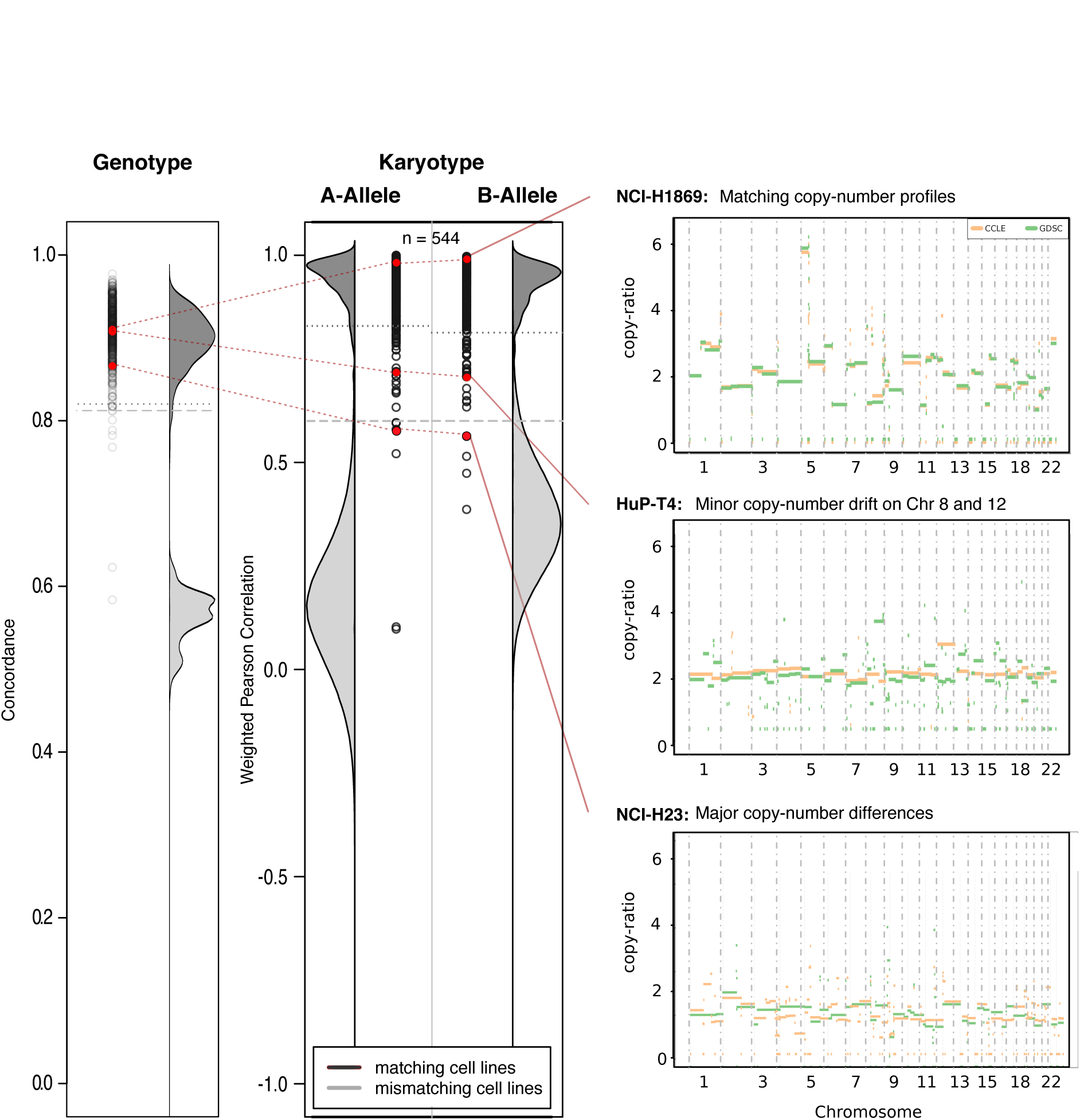
Weighted Pearson correlation for homologue-specific copy ratios of cell lines with matching annotations between GDSC and CCLE. Each cell line of interest has their corresponding genotype concordance marked to illustrate comparable genotypes but discordant karyotypes. Density distributions are generated for all cell line pairs with the same annotation (dark grey) and all cell line pairs with different annotations (light grey). The correlation between matching pairs can be used as a surrogate measure of genetic drift as illustrated by NCI-H1869, HuP-T4 and NCI-H23.

### Phenotypic consequence of genetic drift

To test for phenotypic consequences of discordant karyotypes in CCLE and GDSC, we compared genomic features of 397 cell lines against gene expression and drug sensitivity profiles of 14 compounds. By comparing gene expressions between GDSC and CCLE, we observed high correlation between homonymous cell line pairs (*matching*: r=0.81±0.05) and low correlation for non-homonymous (*non-matching*: r=0.58±0.07). When comparing drug sensitivity profiles between these datasets, we calculated the mean ABC of homonymous pairs (*matching*) as 0.09±0.09 and 0.29±0.16 for non-homonymous (*non-matching*), with significant separation per curve (Kolmogorov-Smirnov test p<0.0001). We found that paclitaxel had the largest overlap between matching and non-matching groups but still remained discriminative (Kolmogorov-Smirnov test p 5×10^−4^; Supp. Fig. 4).

We examined two homonymous cell line pairs in GDSC and CCLE with the lowest concordance between karyotypes for disparities in their phenotypes. The acute lymphocytic leukemia cell line REH presented with a mostly diploid genome with few large aberrations (r_HSCR-A_=0.519, r_HSCR-B_=0.677) (Supp. Fig. 5a), while the lung adenocarcinoma cell line NCI-H23 exhibited a more aneuploid profile with a high number of CNV differences (r_HSCR-A_=0.578, r_HSCR-B_=0.562) (Supp. Fig. 5b). A comparison of gene expression profiles revealed correlations on the lower spectrum of the *matching* distribution (z_REH_=-1.71, z_NCI-H23_=-0.94), but were still more similar when compared to the *non-matching* distribution (likelihood ratio LR_REH_=-3.72, LR_NCI-H23_=-7.13). When comparing drug sensitivity profiles, we found that REH and NCI-H23 were slightly more likely to be associated with a discordant drug sensitivity profile for paclitaxel (LR_REH_=0.52, LR_NCI-H23_=0.52) and PD-0332991 (LR_REH_=1.16, LR_NCI-H23_=1.11). REH was also more discordant for lapatinib (LR=1.25) and PHA-665752 (LR=2.02), however, the AAC values were low and may just represent variance in the drug sensitivity assay (Fig. 5)^6,21,30^. These results suggests that the discordant karyotypes may be associated with differential drug sensitivity while retaining a comparable transcriptome.

**Figure 5.**
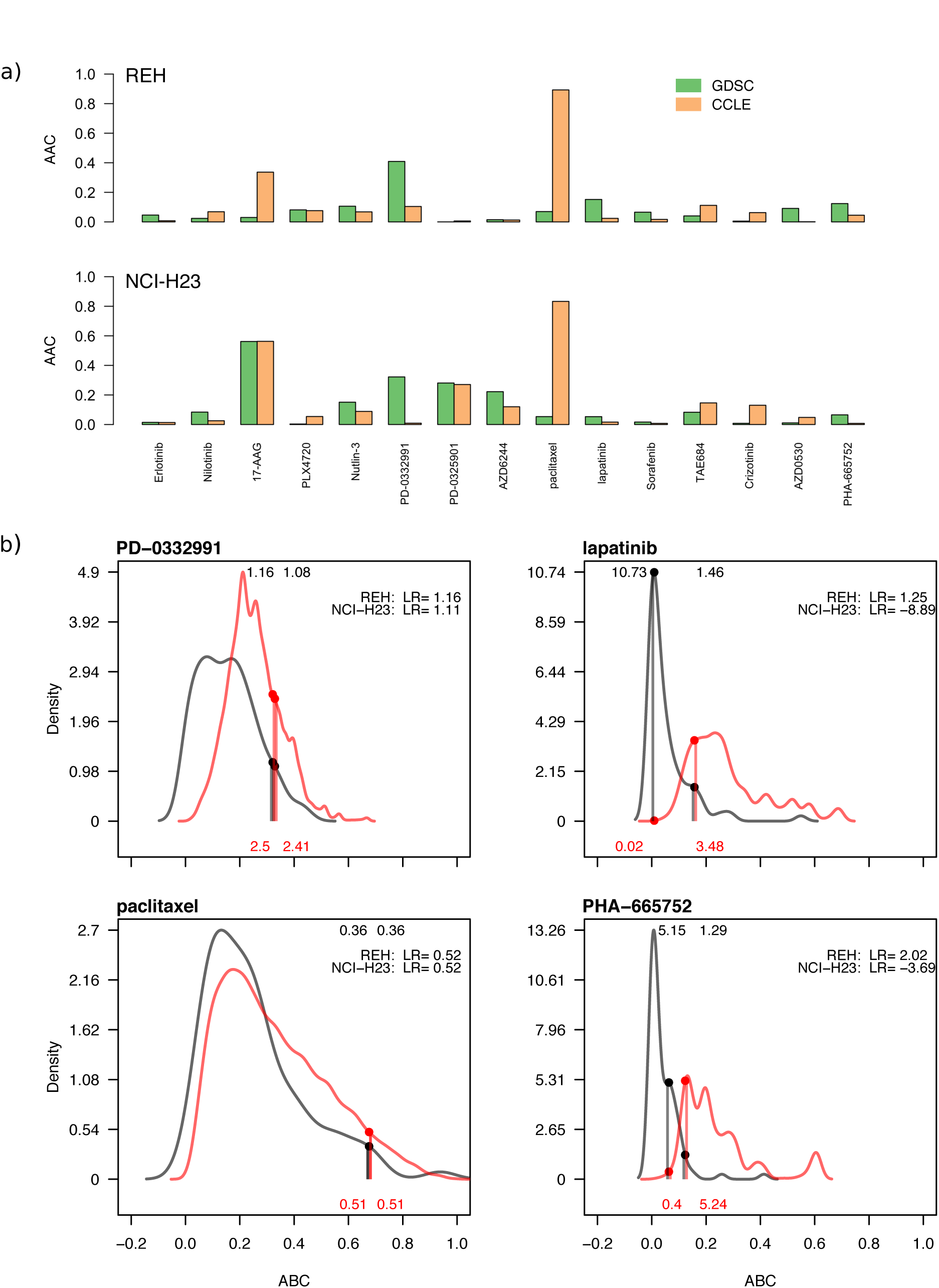
Drug sensitivity profiles for the cell lines REH and NCI-H23, exemplar cases of genetic drift between the GDSC and CCLE dataset. a) The area above the curve (AAC) of the dose-response curves for each overlapping drugs tested on both cell lines in both datasets. b) Area between the density curves between matching cell lines (black) and non-matching cell lines (red). The likelihood values of NCI-H23 and REH are reported for each curve, as well as the likelihood ratio (LR) of non-matching to matching likelihoods.

To evaluate whether these disparities in drug sensitivity profiles can be explained by CNV differences in REH and NCI-H23, we referenced the online LOBICO database ^31^, a tool designed to predict multivariate features influencing drug sensitivity. A predictive feature of PD-0332991 was the absence of genes *FAT1* and *IRF2* due to a 4q deletion, and absence of a 3p14.2 deletion, both conditions that were violated in NCI-H23 screened by CCLE. These predictions corroborate the drug sensitivity profiles displaying CCLE’s NCI-H23 as more resistant (AAC=0.009) when compared to GDSC (AAC=0.321). Similarly, predictive features for paclitaxel sensitivity were a *MLL2* mutation and deletion of 16q23.1 and 18q22.1. CCLE’s NCI-H23 harbored a deletion of both 16q23.1 and gain of 18q22.1 relative to GDSC which may be related to increased paclitaxel sensitivity (AAC_CCLE_=0.89, AAC_GDSC_=0.069; Fig. 6). While REH harbored no CNV on any of the predictive features of PD-0332991, this line did contain an amplification of chromosome 16 in GDSC relative to CCLE that may be related to the increased resistance to paclitaxel (AAC_GDSC_=0.069, AAC_CCLE_=0.892; Supp. Fig. 6). These results exemplify scenarios where CNV-discrepancies in isogenic cell lines may be a driving force for differential phenotypes, however, the inherent noise in drug sensitivity profiles obscures direct links between phenotype and CNV disparities.

**Figure 6.**
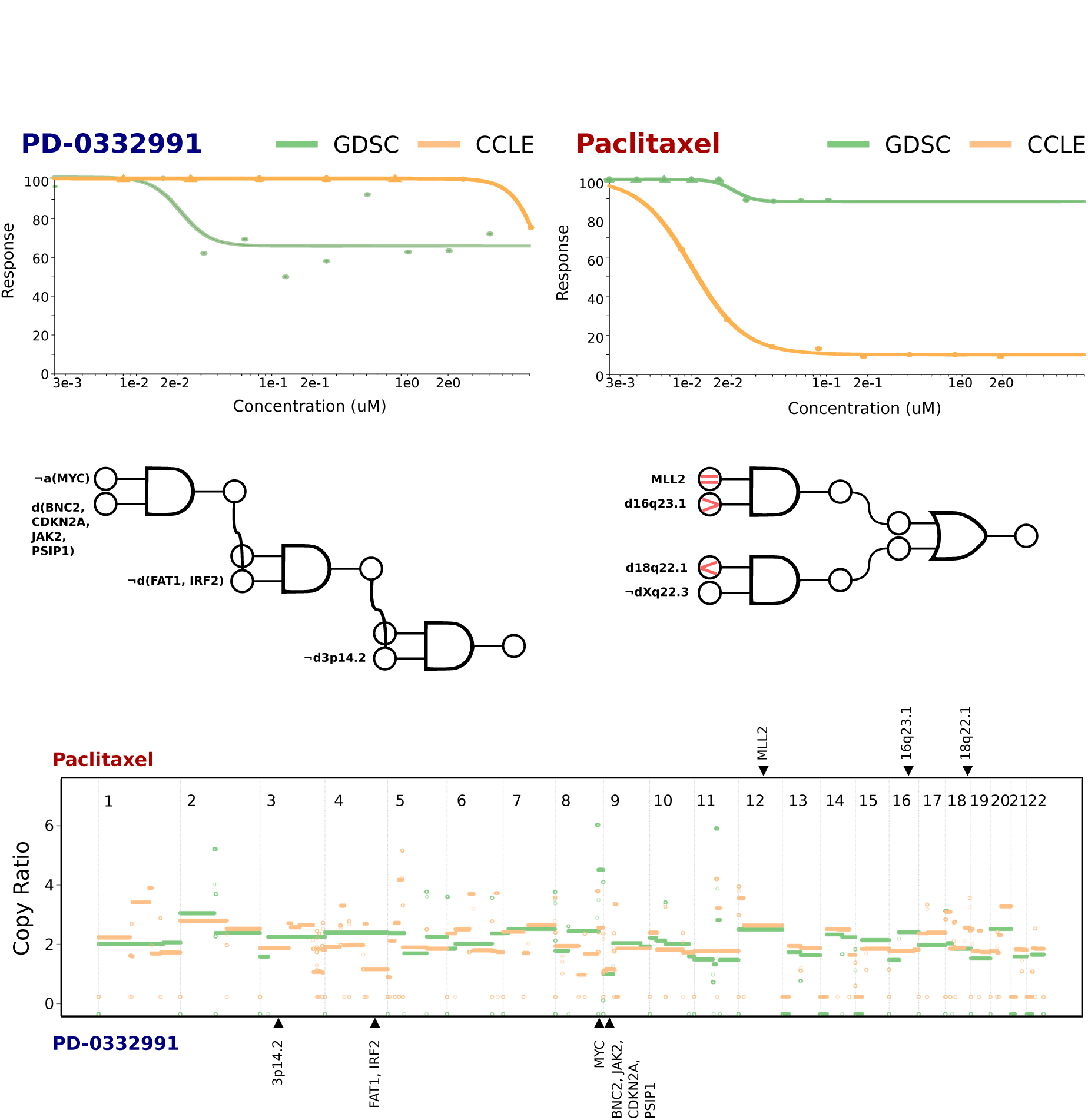
A comparison of drug sensitivity profiles for the lung adenocarcinoma cell line NCI-H23 between the GDSC and CCLE datasets. The top panels illustrate divergent drug-sensitivity profiles for PD-0332991, a CDK4/6 inhibitor, and paclitaxel, a microtubule stabilizing chemotherapeutic. The middle panels illustrate top predictive models for each drug, pre-calculated by the Netherlands Cancer Institute using their LOBICO algorithm. Grey “>”, “<”, and “=” symbols are used to indicate instances where the GDSC line has a CNV greater than, less than, or equal to the CNV state in CCLE. The bottom panel is the GDSC and CCLE copy-number profiles quantile-normalized to make them more comparable; Tracks above and below the CNV figure corresponds to the LOBICO features for paclitaxel or PD-0332991.

### Comparison with Illumina Omni 2.5M Platform

To illustrate how an external dataset can be projected against those already analyzed, we compared all cell line molecular profiles against a study by the Genentech Cell Line Screening Initiative (gCSI) ^7^ which ran 668 cancer cell lines on the Illumina Omni 2.5M SNP-microarray platform. A cross-comparison of genotypes from gCSI to all reference datasets (CCLE, GDSC, CGP, and Pfizer) using 337,353 shared SNPs authenticated all cell line identities. Using these data, we generated usable copy-number profiles for 644/668 cell lines, the majority of which were concordant allele-specific copy-number profiles in our reference datasets (r_A-allele_ = 0.79±0.07, r_B-allele_ = 0.85±0.08). Breaking down this distribution for the A and B alleles, 210 and 362 cell lines with *matching* concordance (r > 0.8), 190 and 79 that were *ambiguous* (0.6 > r > 0.8), and 3 and 4 that were *non-matching* (r < 0.6), respectively. Overall, we observed similar trends of well-controlled isogenic cell lines with karyotypes that ranged on a continuous spectrum of discordance (Supp. Fig. 7).

To compare phenotypic measures, we conducted a comparison of expression and drug sensitivity profiles of 185 gCSI cell lines across 6 compounds against the CCLE dataset as both studies used similar drug sensitivity assays ^6^. Overall, we showed that cell lines with CNV discordance (ambiguous) have similar gene expression profiles (match), albeit on the lower end of the correlation range (LR_Match_= −7.7±0.5; LR_Ambiguous_= −7.4±0.3; LR_Non-match_= −5.8±2.0); a comparison of drug sensitivity profiles also did not show discordances proportional to the severity of CNVs (Fig. 7). While we measured a significant difference between all *matching* and *non-matching* drug sensitivity distributions (Kolmogorov-Smirnov test p<0.0001), there was a large amount of overlap between these distributions for paclitaxel and irinotecan which limited the range of discrepant values to the extremes. Overall, most cell line pairs had drug sensitivity profiles that were better explained by the *matching* distribution except for PSN-1, KYSE-510, and KARPAS-620 being more discordant for PD-0325901 (LR=1.60, LR=0.3, LR=0.1) and NUGC-3 being more discordant for erlotinib (LR=2.0) (Fig. 7).

**Figure 7.**
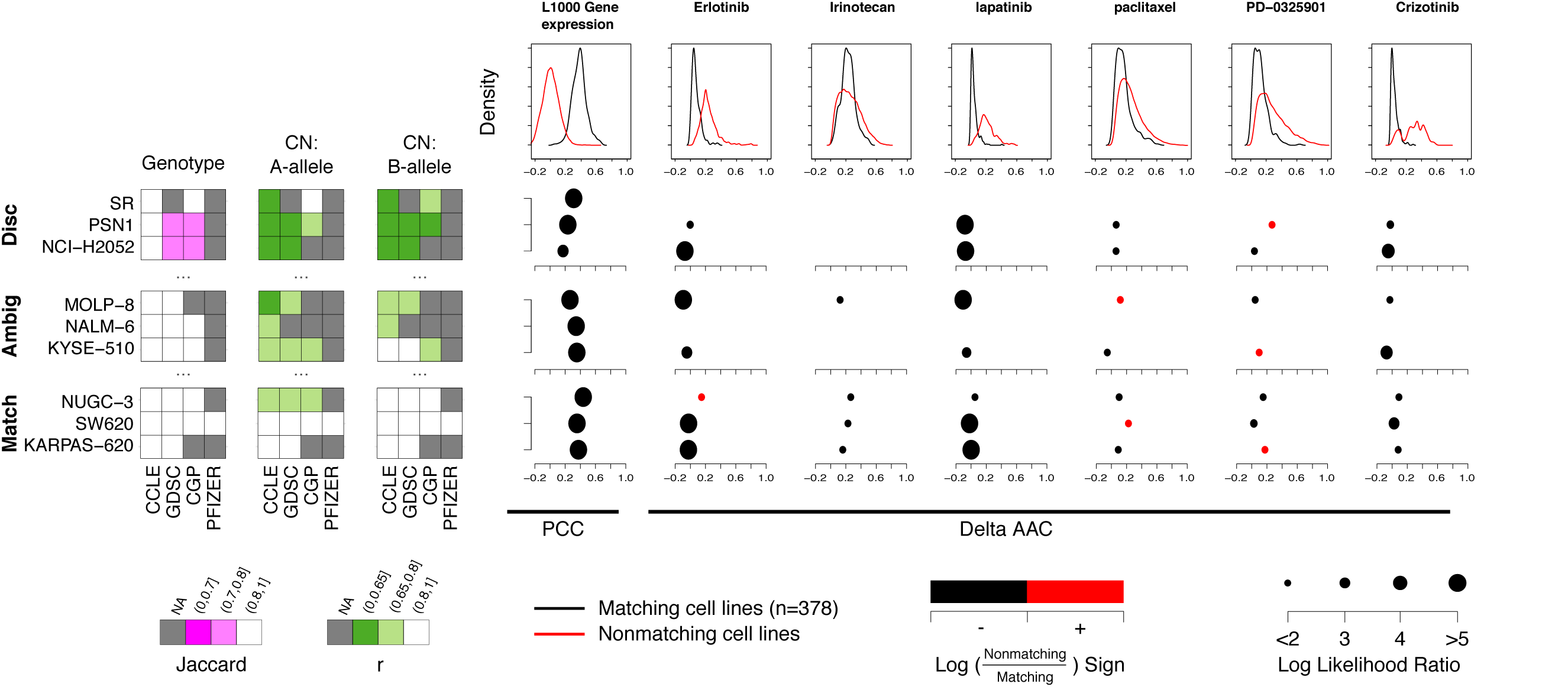
Example comparisons of genotype and copy-number profiles from gCSI cell lines to the reference CCLE, GDSC, CGP, and Pfizer datasets for cell lines with discordant (Disc) CNV profiles, ambiguous (Ambig) CNV profiles, and matching (Match) CNV profiles. Phenotype comparisons are made between 378 overlapping cell lines between the gCSI and CCLE datasets using gene-expression profiles subsetted for the L1000 geneset, as well as the area between the dose-response curves (ABC) of 6 overlapping drugs used in both studies. Log likelihood ratios for cell lines with discordant phenotypes over concordant phenotypes are represented as a dot plot.

As PSN-1 had highly dissimilar karyotypes between gCSI and CCLE paired with substantially different response to the MEK-inhibitor PD-0325901, we investigated whether there were informative CNVs that could explain the difference in drug response. The most predictive LOBICO model was based on oncogenic mutations in *BRAF*, *KRAS* or *NRAS*, which are located at 7q34, 12p12.1, and 1p13.2 respectively. We observed an amplification of 7q34 (containing *BRAF*) in gCSI relative to CCLE and equivalent copy-numbers of 12p12.1 (containing *KRAS*) and 1p13.2 (containing *NRAS*). The amplification of *BRAF* in gCSI was concordant with increased sensitivity relative to CCLE (Supp. Fig. 8). While we show that CNV discordance in PSN-1 may be associated with drug sensitivity, we did not see any overlap between the predictive features with KYSE-510 and KARPAS-620, possibly due to the limited number of CNV discordances in these two cell lines. Similarly, we saw no overlap of predictive features of erlotinib sensitivity with NUGC-3, again possibly due to the limited number of CNV discordances and the highly targeted nature of erlotinib.

## DISCUSSION

Our investigation of the six largest pharmacogenomic studies published to date revealed high-quality datasets with few genomic discrepancies. While we expected that ∼18% of cell lines would be misidentified per dataset ^32^, we only detected only 6 out of 1,378 cell lines (0.4%) discordant in genetic and karyotypic identity in multiple datasets; highlighting the rigorous experimental protocols used in large-scale pharmacogenomic studies. We nominated an additional 51 cell lines with discrepant genotypes (3.7%), although the majority of these were marginally discordant. Importantly, we detected 311 heteronymous but isogenic cell lines; 186 of which likely originated from the same patient and 94 which were cases of misidentification/contamination.

Standard cell line authentication by profiling a small number of short-tandem repeats (STR) is insufficient to describe genomic heterogeneity within isogenic cell lines. Herein, we report on 47 isogenic cell line pairs across all datasets with poor karyotype concordance. This heterogeneity as measured by CNV discrepancies may be a factor that results with disagreement between drug sensitivity measurements across pharmacogenomic datasets ^6,19,20,22^, however, the inherent noise in drug sensitivity profiles makes it difficult to attribute drift to phenotype discordances. The use of predictive features of drug sensitivity as calculated by an external group ^31^ indicate that the CNV discordances in the cell lines analyzed could have a predictable effect on drug response.

While a differential pharmacological response to molecular variation is a tenet of personalized medicine, Ben-David et al. has shown that 82% of differentially active compounds in 27 strains of the MCF7 cancer cell line can be explained by their gene expression signatures ^18^. These differential responses extend further than just a discrepant transcriptome. For example, Kleensang et al. has shown that karyotype discordances in MCF-7 cell lines affects response to estrogen agonists ^33^, similar to our finding of CNV discordances associating with discrepant pharmacological responses. Not only can response to therapeutic compounds be ploidy-dependent ^34–36^, but CNVs can also be rapidly acquired due to selective pressures in a few cell cycles ^37,38^. For instance, gain of isochromosome 5L in *C.albicans* is rapidly acquired to confer resistance to fluconazole ^39^. Thus, the innate instability and heterogeneity of cancer cell lines is an essential factor to consider when considering pharmacological assays, especially as external pressures can exacerbate inherent heterogeneity.

In conclusion, our study highlights that genomic and chromosomal instability is an underestimated confounder that can result in differential pharmacological response in isogenic cell lines. While our study is limited by limited detection of smaller CNVs in the absence of normalization, we demonstrate that large-scale events have phenotypic effects and it is reasonable to expect smaller, focal events may also have an effect. We also have limited confidence to classify drug sensitivity discordance due to large variability in drug sensitivity measurements for homonymous cell lines. To enable high confidence associations between heterogeneity and drug response, we would require a well-controlled experiment between two institutes similar to Kleensang et al. ^33^, or repeat drug sensitivity assays on single-cell cultures similar to Ben-David et al. ^18^. Even so, our analysis shows the power of integrated pharmacogenetics data sets to cross-validate drug sensitivity results and to further refine mechanisms underpinning these associations. The commitment to open science of the investigators participating in the six contributing studies is laudable as we will continue to derive additional insight as we augment this resource with additional data.

