## Supplemental tables and figures for "Karyotypic divergence confounds cellular phenotypes in large pharmacogenomic studies"

| Cell-line 1 | Cell-line 2 | ICLAC-1 | ICLAC-2 | Classification | CCLE Status |
|---|---|---|---|---|---|
| AU565 | SK-BR-3 | No | No | match | Metastatic pair: SK-BR-3 and AU565 are from the same patient |
| C3A | Hep G2 | No | No | match | Identical lines: Hep G2 and C3A share high SNP identity |
| CMK | CMK-11-5 | No | No | match | Identical lines: CMK, CMK-86 and CMK-11-5 share high SNP identity |
| CMK | CMK-86 | No | No | match | Identical lines: CMK, CMK-86 and CMK-11-5 share high SNP identity |
| FTC-133 | FTC-238 | No | No | match | Metastatic pair: FTC-133 and FTC-238 are from the same patient. FTC-133 was derived from the primary tumor. |
| DLD-1 | HCT-15 | No | No | match | Identical lines: HRT18, HCT15 and DLD1 share high SNP identity |
| DLD-1 | HRT-18 | No | No | mismatch | Identical lines: HRT18, HCT15 and DLD1 share high SNP identity |
| HCT-15 | HRT-18 | Yes | No | mismatch | Identical lines: HRT18, HCT15 and DLD1 share high SNP identity |
| HLE | HLF | No | No | match | Identical lines: HLE and HLF share high SNP identity |
| 143B | HOS | No | No | match | Identical lines: HTK-, HOS and 143B share high SNP identity and are very likely to be osteosarcoma |
| HOS | HTK- | No | No | mismatch | Identical lines: HTK-, HOS and 143B share high SNP identity and are very likely to be osteosarcoma |
| 143B | HTK- | No | No | match | Identical lines: HTK-, HOS and 143B share high SNP identity and are very likely to be osteosarcoma |
| IGR-37 | IGR-39 | No | No | match | Metastatic pair: IGR-37 and IGR-39 are from the same patient. IGR-37 is a metastasis |
| MONO-MAC-1 | MONO-MAC-6 | No | No | match | MONO-MAC-1 and MONO-MAC-6 were derived from the same patient |
| PA-TU-8988S | PA-TU-8988T | No | No | match | Identical lines: PA-TU-8988T and PA-TU-8988S are from the same patient |
| PANC-10-05 | PL45 | No | No | match | Identical lines: Panc 10.05 and PL45 were derived from the same sample |
| RH-30 | SJRH30 | No | No | match | Identical lines: SJRH30 and RH30 share high SNP identity |
| SH-SY5Y | SK-N-SH | No | No | match | Identical lines: SH-SY5Y is a subclone derived from SK-N-SH and is identical to LAN-5 |
| 1321N1 | U-118-MG | No | No | match | Identical lines: U-118 MG, U-138 MG and 1321N1 share high SNP identity |
| U-118-MG | U-138 MG | Yes | Yes | mismatch | Identical lines: U-118 MG, U-138 MG and 1321N1 share high SNP identity |
| 1321N1 | U-138 MG | No | Yes | mismatch | Identical lines: U-118 MG, U-138 MG and 1321N1 share high SNP identity |
| WM-115 | WM-266-4 | No | No | match | Metastatic pair: WM-115 and WM-266-4 are from the same patient |
| LC-1/sq-SF | LC-1F | No | No | match | Identical lines: LC-1/sq-SF and LC-1F share high SNP identity. |
| MEC-1 | MEC-2 | No | No | match | MEC-1 and MEC-2 were derived from the same patient |
| CGTH-W-1 | SW579 | No | No | mismatch | Identical lines: SW579 and CGTH-W-1 share high SNP identity. CGTH-W-1 really is SW579. |
| COLO-800 | COLO-818 | Yes | Yes | mismatch | Identical lines: COLO-818 and COLO-800 share high SNP identity and are likely to be COLO-800 |
| COLO-829 | COLO-849 | No | No | mismatch | Identical lines: COLO 829 and COLO-849 share high SNP identity |
| HPAC | KCI-MOH1 | Yes | Yes | mismatch | Identical lines: KCI-MOH1 is a derivative of HPAC |
| AZ-521 | HUTU-80 | Yes | Yes | mismatch | Identical lines: HuTu 80 and AZ-521 share high SNP identity and are likely to be HuTu 80 |
| KARPAS-422 | OCI-LY10 | No | No | mismatch | Identical lines: KARPAS-422 and OCI-LY10 share high SNP identity. |
| KPL-1 | MCF7 | Yes | Yes | mismatch | Identical lines: MCF-7 and KPL-1 share high SNP identity |
| OC 316 | OC-314 | No | No | mismatch | Identical lines: OC314, OC315 and OC316 share high SNP identity |
| ONCO-DG-1 | OVCAR-3 | Yes | Yes | mismatch | Identical lines: NIH:OVCAR-3 and ONCO-DG-1 share high SNP identity. Likely to be NIH:OVCAR-3 |
| GR-M | PSN1 | No | No | mismatch | Identical lines: GR-M and PSN1 share high SNP identity and are likely to be really PSN1 (pancreas) |
| COLO 775 | RPMI-8226 | No | Yes | mismatch | Identical lines: RPMI 8226, COLO 775 and COLO-677 share high SNP identity and are likely to be RPMI 8226 |
| SW480 | SW620 | Yes | Yes | mismatch | Metastatic pair: SW620 and SW480 were derived from the same patient |
| Hs 571.T | TOV-112D | No | No | mismatch | Identical lines: TOV-112D and Hs 571.T share high SNP identity |
| SNB19 | U251 | Yes | Yes | mismatch | Identical lines: SNB-19 and U-251 MG share high SNP identity |
| HLC-1 | HMC-1-8 | No | No | mismatch | Identical lines: HMC-1-8 and HLC-1 share high SNP identity and are probably lung. |
| YMB-1 | ZR751 | Yes | Yes | mismatch | Identical lines: ZR-75-1 and YMB-1 share high SNP identity |
| KLM-1 | PK-1 | No | No | mismatch | PK-1 is derived from the same patient as KLM-1 cells |
| KE-97 | KMS-18 | No | No | mismatch | Identical lines: KMS-18 and KE-97 share high SNP identity. |
| EB1 | EB2 | No | No | possible match | Identical lines: EB1 and EB2 share high SNP identity |
| HEL | HEL 92.1.7 | No | No | possible match | Identical lines: HEL 92.1.7 is a subclone of HEL |
| Alexander cells | PLC-PRF-5 | No | No | possible match | Identical lines: Alexander cells and PLC/PRF/5 share high SNP identity |
| CMK-11-5 | CMK-86 | No | No | possible match | Identical lines: CMK, CMK-86 and CMK-11-5 share high SNP identity |
| HEC-1 | HEC-1-A | No | No | match | Identical lines: HEC-1-A HEC-1-B are subclones of HEC-1 and share high SNP identity |
| RT-112 | RT112/84 | No | No | match | RT112/84 is a clonal derivative of RT112 |

Supplementary Table 1: Cell lines pairs that have matching genotypes within the CCLE dataset paired with CCLE-descriptions for these discrepancies.

| Dataset 1 | Dataset 2 | Discordant CNV Profiles | Discordant SNP Profiles | Intersect |
|---|---|---|---|---|
| CCLE | CGP | AN3-CA, HT-29, HUH-6-clone5, HuP-T4, KNS-81-FD, LOXIMVI, MCF7, MDA-MB-157, MDA-MB-415, MDA-MB-435, Mewo, MOG-G-CCM, NB1, NCI-H2087, NCI-H2122, NCI-H23, OVCAR-3, PC-3, PF-382, REH, SCC-15, SK-MEL-24, SR, U251, UACC-893 | EPLC-272H1, HCC19371, HuH-7, KP-41, KYSE-701, LC-1F, MDA-MB-4681, MOG-G-CCM1, VM-CUB-11 | COR-L51, NB4, SW403 |
| CCLE | Pfizer | CAKI-1, HCC38, LOXIMVI, MDA-MB-435, U251 | BT-20, NCI-H1184 | |
| CCLE | Pfizer2 | HCC38, UACC-812 | BT-20 | |
| CCLE | GDSC | DOHH-2, HCC-33, HCC-78, Hs 939.T, Hs 940.T, HT-29, HuP-T4, JHOS-2, JHOS-4, KNS-81-FD, LOXIMVI, MCF7, MDA-MB-157, MDA-MB-415, Mewo, NAMALWA, NB1, NCI-H1568, NCI-H2009, NCI-H2066, NCI-H2087, NCI-H2122, NCI-H23, OCI-AML5, OE21, OUMS-23, OVCAR-3, PC-3, REH, SCaBER, SCC-15, Set-2, SK-MEL-24, U-2-OS, U251, UACC-812, UACC-893 | EPLC-272H, FU97, HCC19371, HuCCT1, KP-4, KYSE-70, MDA-MB-4361, MDA-MB-4681, NCI-H211, TT2, VM-CUB-1 | COLO-783, Ishikawa (Heraklio) 02 ER-, MOG-G-CCM |
| CCLE | GDSC2 | KM-H2 | | |
| CGP | Pfizer | CAKI-1, EKVX, HCT-15, MCF7, MDA-MB-157, MDA-MB-415, NCI-H23, NCI-H522, OVCAR-3, PC-3, SNB75 | HCC19371, MDA-MB-4681, OVCAR-51, UACC-8121 | |
| CGP | Pfizer2 | BT-549, MCF7, MDA-MB-157, MDA-MB-415, UACC-812 | | |
| CGP | GDSC | | A253, CAL-12T, COLO-829, DMS-114, HuH-7, IA-LM, LB2241-RCC, LCLC-103H, NCI-H1299, SCC-15, SNU-423, TT1, U-2-OS, UACC-8121, UACC-893, VA-ES-BJ | HUH-6-clone5, NB4, SW403 |
| CGP | GDSC2 | KM-H2 | | |
| Pfizer | Pfizer2 | | SUM185PE, UACC-812 | |
| GDSC | Pfizer | CAKI-1, EKVX, HCC1937, HCT-15, MCF7, MDA-MB-157, MDA-MB-415, NCI-H23, NCI-H522, OVCAR-3, PC-3, SNB75, UACC-812 | HCC19372, MDA-MB-4362, MDA-MB-4682, OVCAR-5, UACC-8122 | |
| GDSC | Pfizer2 | BT-549, MCF7, MDA-MB-157, MDA-MB-415, UACC-812 | MDA-MB-4363, MDA-MB-4683 | |
| GDSC | Pfizer3 | BT-549 | MDA-MB-4364 | |
| GDSC | GDSC2 | KM-H2 | | |

Supplementary Table 2: Cell lines between two datasets with discordant CNV or SNP profiles as well as the cell lines that are discordant in CNV and SNP.

| Cell Line | Dataset | Tissue | Morphology | Age | Sex | Ethnicity | Cell Line | Dataset | Tissue | Morphology | Age | Sex | Ethnicity |
|---|---|---|---|---|---|---|---|---|---|---|---|---|---|
| AU565 | CGP | Breast | Carcinoma | 32 | F | Caucasian | SK-BR-3 | CCLE | Breast | Carcinoma | 32 | F | Caucasian |
| AU565 | CCLE | Breast | Carcinoma | 32 | F | Caucasian | SK-BR-3 | CCLE | Breast | Carcinoma | 32 | F | Caucasian |
| AU565 | Pfizer | Breast | Carcinoma | 32 | F | Caucasian | SK-BR-3 | CCLE | Breast | Carcinoma | 32 | F | Caucasian |
| AU565 | Pfizer | Breast | Carcinoma | 32 | F | Caucasian | SK-BR-3 | CCLE | Breast | Carcinoma | 32 | F | Caucasian |
| AU565 | GDSC | Breast | Carcinoma | 32 | F | Caucasian | SK-BR-3 | CCLE | Breast | Carcinoma | 32 | F | Caucasian |
| BB30-HNC | CGP | | HnC Carcinoma | | F | | BB30-PBL | CGP | PBL | HnC Carcinoma | | F | |
| BB30-HNC | GDSC | | HnC Carcinoma | | F | | BB30-PBL | CGP | PBL | HnC Carcinoma | | F | |
| BB49-HNC | CGP | | HnC Carcinoma | | F | | BB49-EBV | CGP | | HnC Carcinoma | | F | |
| BB49-HNC | GDSC | | HnC Carcinoma | | F | | BB49-EBV | CGP | | HnC Carcinoma | | F | |
| BB65-RCC | CGP | | RCC | | M | | BB65-EBV | CGP | | RCC | | M | |
| BB65-RCC | GDSC | | RCC | | M | | BB65-EBV | CGP | | RCC | | M | |
| BE2-M17 | GDSC | Brain | NB | 2 | M | | SK-N-BE(2) | CCLE | Brain | NB | 2 | M | |
| C2BBe1 | CGP | Colon | CRAC | 72 | M | Caucasian | CACO2 | Pfizer | Colon | CRAC | 72 | M | Caucasian |
| C2BBe1 | CCLE | Colon | CRAC | 72 | M | Caucasian | CACO2 | Pfizer | Colon | CRAC | 72 | M | Caucasian |
| C2BBe1 | GDSC | Colon | CRAC | 72 | M | Caucasian | CACO2 | Pfizer | Colon | CRAC | 72 | M | Caucasian |
| C3A | CCLE | Liver | HCC | 15 | M | Caucasian | Hep G2 | CCLE | Liver | HCC | 15 | M | Caucasian |
| C3A | GDSC | Liver | HCC | 15 | M | Caucasian | Hep G2 | CCLE | Liver | HCC | 15 | M | Caucasian |
| C4I | GDSC | Cervix | Carcinoma | 41 | F | Caucasian | C-4-II | CGP | Cervix | Carcinoma | 41 | F | Caucasian |
| CMK | CGP | | AMegL | 10mo | M | | CMK-11-5 | CCLE | | AMKL | | | |
| CMK | CCLE | | AMegL | 10mo | M | | CMK-11-5 | CCLE | | AMKL | | | |
| CMK | GDSC | | AMegL | 10mo | M | | CMK-11-5 | CCLE | | AMKL | | | |
| CMK | CGP | | AMegL | 10mo | M | | CMK-86 | CCLE | | AMegL | | | |
| CMK | CCLE | | AMegL | 10mo | M | | CMK-86 | CCLE | | AMegL | | | |
| CMK | GDSC | | AMegL | 10mo | M | | CMK-86 | CCLE | | AMegL | | | |
| COLO-829 | CGP | Skin | MM | 45 | M | Caucasian | COLO-829-BL | CGP | Skin | MM | 45 | M | Caucasian |
| COLO-829 | CCLE | Skin | MM | 45 | M | Caucasian | COLO-829-BL | CGP | Skin | MM | 45 | M | Caucasian |
| COLO-829 | GDSC | Skin | MM | 45 | M | Caucasian | COLO-829-BL | CGP | Skin | MM | 45 | M | Caucasian |
| EoL-1-cell | CGP | | | | | | EOL-1 | CCLE | | | | | |
| EoL-1-cell | GDSC | | | | | | EOL-1 | CCLE | | | | | |
| FTC-133 | CCLE | | FTC | 42 | M | | FTC-238 | CCLE | | FTC | 42 | M | |
| GA-10 | CCLE | | | | | | GA-10-Clone-4 | CGP | | | | | |
| GA-10 | GDSC | | | | | | GA-10-Clone-4 | CGP | | | | | |
| GP5d | CGP | | CRC | 71 | F | | Gp2D | CCLE | | CRC | 71 | F | |
| GP5d | GDSC | | CRC | 71 | F | | Gp2D | CCLE | | CRC | 71 | F | |
| H3255 | GDSC | | | | | | NCI-H3255 | CCLE | | | | | |
| HCC1599 | CGP | | | | | | HCC1599-BL | CGP | | | | | |
| HCC1599 | Pfizer | | | | | | HCC1599-BL | CGP | | | | | |
| HCC1599 | Pfizer | | | | | | HCC1599-BL | CGP | | | | | |
| HCC1599 | GDSC | | | | | | HCC1599-BL | CGP | | | | | |
| HCC1937 | CGP | | | | | | HCC1937-BL | CGP | | | | | |
| HCC1937 | GDSC | | | | | | HCC1937-BL | CGP | | | | | |
| HCC1954 | CGP | | | | | | HCC1954-BL | CGP | | | | | |
| HCC1954 | CCLE | | | | | | HCC1954-BL | CGP | | | | | |
| HCC1954 | Pfizer | | | | | | HCC1954-BL | CGP | | | | | |
| HCC1954 | Pfizer | | | | | | HCC1954-BL | CGP | | | | | |
| HCC1954 | GDSC | | | | | | HCC1954-BL | CGP | | | | | |
| HCC2157 | CGP | | | | | | HCC2157-BL | CGP | | | | | |
| HCC2218 | CGP | | | | | | HCC2218-BL | CGP | | | | | |
| HCC2218 | CCLE | | | | | | HCC2218-BL | CGP | | | | | |
| HCC2218 | Pfizer | | | | | | HCC2218-BL | CGP | | | | | |
| HCC2218 | Pfizer | | | | | | HCC2218-BL | CGP | | | | | |
| HCC2218 | GDSC | | | | | | HCC2218-BL | CGP | | | | | |
| HCC38 | Pfizer | | | | | | HCC38-BL | CGP | | | | | |
| HCT-15 | CGP | Colon | CRAC | | M | | DLD-1 | CCLE | Colon | CRAC | adult | M | |
| HCT-15 | CCLE | Colon | CRAC | | M | | DLD-1 | CCLE | Colon | CRAC | adult | M | |
| HCT-15 | Pfizer | Colon | CRAC | | M | | DLD-1 | CCLE | Colon | CRAC | adult | M | |
| HCT-15 | GDSC | Colon | CRAC | | M | | DLD-1 | CCLE | Colon | CRAC | adult | M | |
| HEC-1 | CGP | | | | | | HEC-1-A | CCLE | | | | | |
| HEC-1 | CCLE | | | | | | HEC-1-A | CCLE | | | | | |
| HEC-1 | GDSC | | | | | | HEC-1-A | CCLE | | | | | |
| HOS | CGP | Bone | OS | 13 | F | Caucasian | 143B | CCLE | Bone | OS | 13 | F | Caucasian |
| HOS | CCLE | Bone | OS | 13 | F | Caucasian | 143B | CCLE | Bone | OS | 13 | F | Caucasian |
| HOS | GDSC | Bone | OS | 13 | F | Caucasian | 143B | CCLE | Bone | OS | 13 | F | Caucasian |
| IGR-37 | CCLE | Lymph | MM | 26 | M | | IGR-39 | CCLE | Skin | MM | 26 | M | |
| IGR-37 | GDSC | Lymph | MM | 26 | M | | IGR-39 | CCLE | Skin | MM | 26 | M | |
| JURKAT | CCLE | PBL | ALL | 14 | M | | J-RT3-T3-5 | CGP | PBL | ALL | 14 | M | |
| JURKAT | GDSC | PBL | ALL | 14 | M | | J-RT3-T3-5 | CGP | PBL | ALL | 14 | M | |
| KMS-12-BM | CGP | | | | | | KMS-12-PE | CGP | | | | | |
| KMS-12-BM | CCLE | | | | | | KMS-12-PE | CGP | | | | | |
| KMS-12-BM | GDSC | | | | | | KMS-12-PE | CGP | | | | | |
| KO52 | GDSC | | | | | | K052 | CCLE | | | | | |
| LC-1-sq | GDSC | | | | | | LC-1/sq-SF | CCLE | Lung | | 69 | M | Japanese |
| LC-1-sq | GDSC | | | | | | LC-1F | CCLE | Lung | | 69 | M | Japanese |
| LC-1/sq-SF | CCLE | Lung | | 69 | M | Japanese | LC-1F | CCLE | Lung | | 69 | M | Japanese |
| LS 180 | CCLE | Colon | CRAC | 58 | F | Caucasian | LS-174T | CGP | Colon | CRAC | 58 | F | Caucasian |
| LS 180 | GDSC | Colon | CRAC | 58 | F | Caucasian | LS-174T | CGP | Colon | CRAC | 58 | F | Caucasian |
| M059J | CGP | Brain | GBM | 33 | M | | M059K | CCLE | Brain | GBM | 33 | M | |
| M059J | GDSC | Brain | GBM | 33 | M | | M059K | CCLE | Brain | GBM | 33 | M | |
| MC-IXC | CGP | Brain | NB | 14 | F | Caucasian | SK-N-MC | CCLE | Brain | NET | 14 | F | Caucasian |
| MC-IXC | GDSC | Brain | NB | 14 | F | Caucasian | SK-N-MC | CCLE | Brain | NET | 14 | F | Caucasian |
| MDA-MB-175-VII | CGP | | | | | | MDAMB175 | Pfizer | | | | | |
| MDA-MB-175-VII | CCLE | | | | | | MDAMB175 | Pfizer | | | | | |
| MDA-MB-175-VII | GDSC | | | | | | MDAMB175 | Pfizer | | | | | |
| MDA-MB-231 | CGP | | | | | | MB231 | Pfizer | | | | | |
| MDA-MB-231 | CCLE | | | | | | MB231 | Pfizer | | | | | |
| MDA-MB-231 | Pfizer | | | | | | MB231 | Pfizer | | | | | |
| MDA-MB-231 | GDSC | | | | | | MB231 | Pfizer | | | | | |
| MEC-2 | CCLE | PBL | CLL | 62 | M | Caucasian | MEC-1 | CCLE | PBL | CLL | 61 | M | Caucasian |
| MONO-MAC-6 | CGP | | AML-M5 | 64 | M | | MONO-MAC-1 | CCLE | | AML-M5 | 64 | M | |
| MONO-MAC-6 | CCLE | | AML-M5 | 64 | M | | MONO-MAC-1 | CCLE | | AML-M5 | 64 | M | |
| MONO-MAC-6 | GDSC | | AML-M5 | 64 | M | | MONO-MAC-1 | CCLE | | AML-M5 | 64 | M | |
| MZ2-MEL | GDSC | | | | | | MZ2-MEL- | CGP | | | | | |

Supplementary Table 3: All cell line pairs across datasets that were discovered to originate from the same patient. REMOVE.

### Supplementary Figures

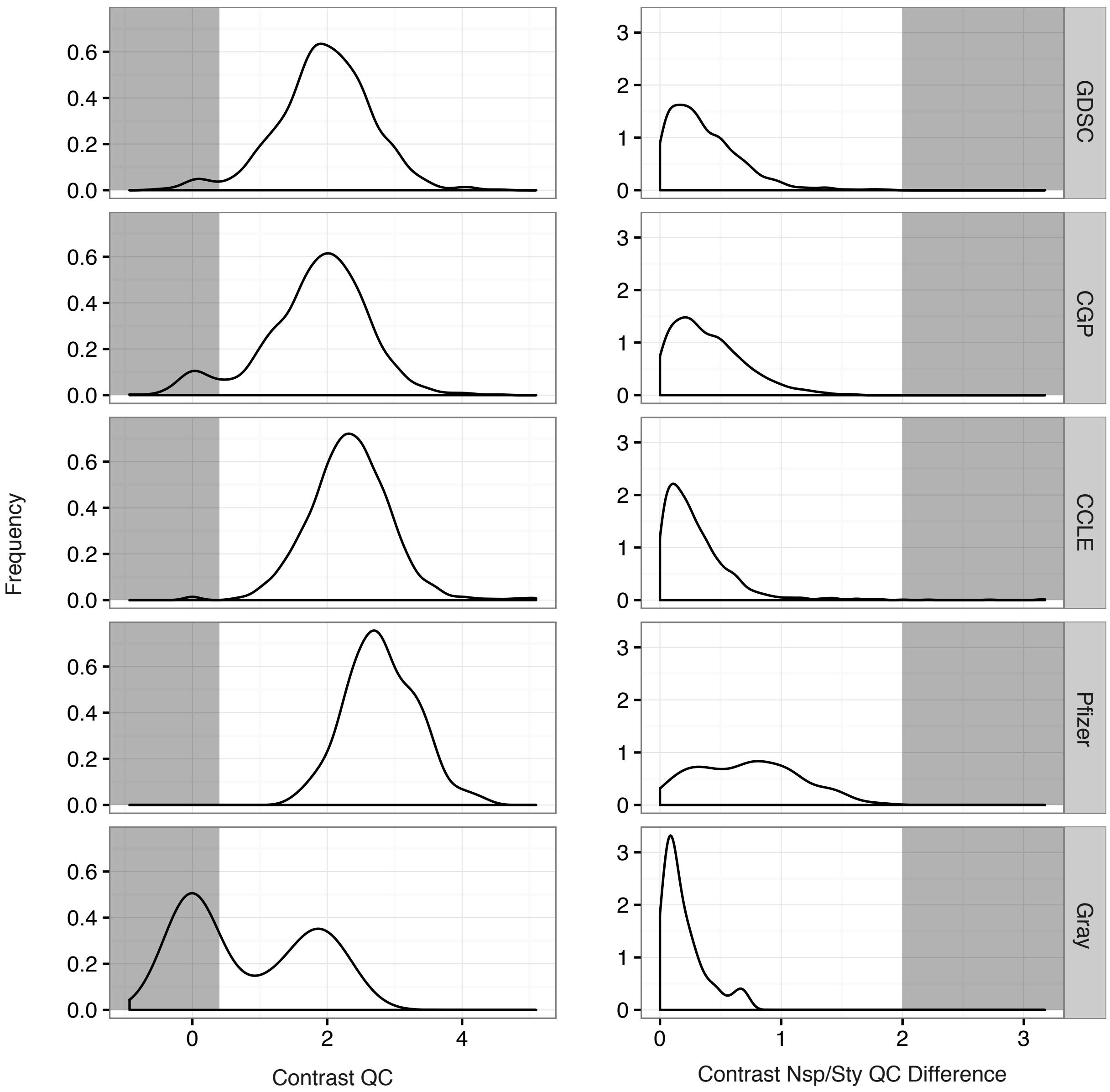


Supplementary Figure 1: Quality control metrics for Affymetrix SNP 6.0 datasets as described by the Affymetrix SNP 6.0 guidelines.

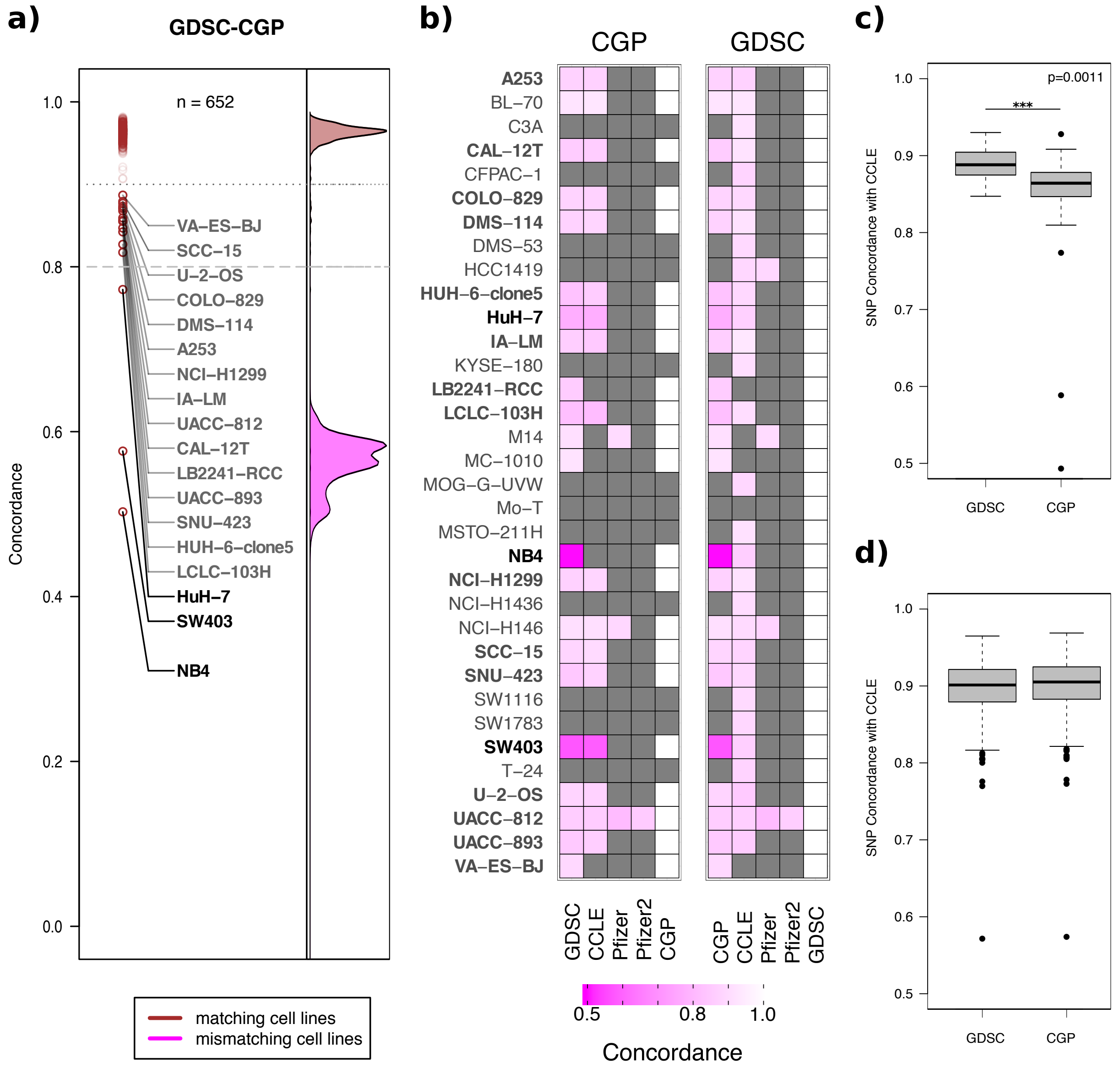


Supplementary Figure 2: A comparison between cell lines from the GDSC and CGP dataset that differ based on their MD5sum digital signature. a) Genotype concordance of cell-line pairs between the GDSC and CGP dataset; two thresholds are shown, the dashed threshold line corresponding to a concordance of 0.8 and the dotted threshold line corresponding to 3 median absolute deviation from the median. b) A heatmap illustrating only cell lines with different digital signatures and how they compare to all other datasets. c) Boxplot showing how the GDSC and CGP cell lines with different digital signatures compare to the CCLE dataset (Wilcoxon rank sum test). d) A negative control to show how GDSC and CGP cell lines with the same digital signatures compare to the CCLE dataset.

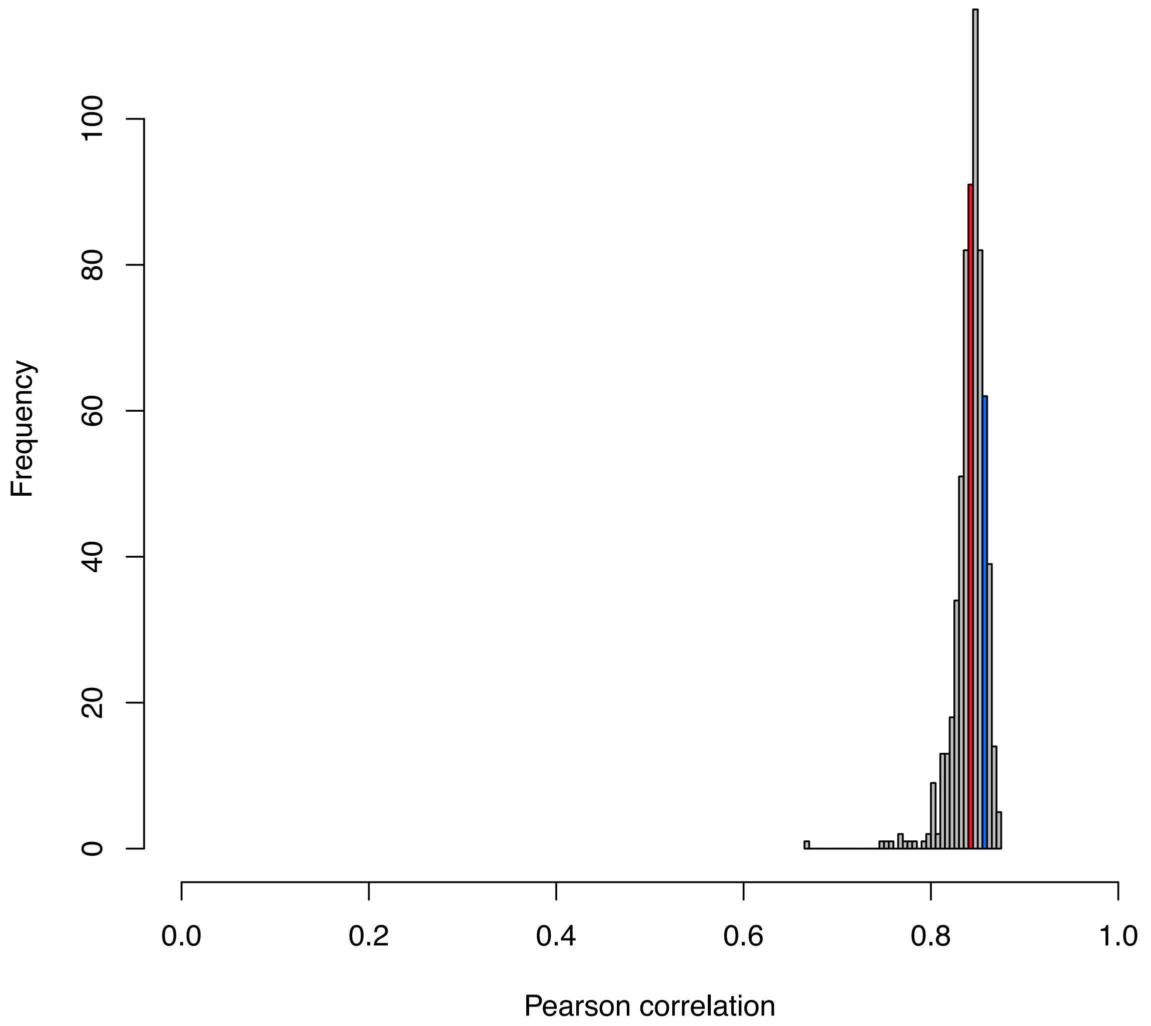


Supplementary Figure 3: Pearson correlation distribution for all cell line pairs with identical annotations between GDSC and CCLE. The red bar corresponds to a Pearson correlation of 0.84 between HPAC screened in GDSC and KCI-MOH1 screened in CCLE. The blue bar corresponds to a Pearson correlation of 0.86 between KARPAS-422 screened in GDSC and OCI-LY10 screened in CCLE.

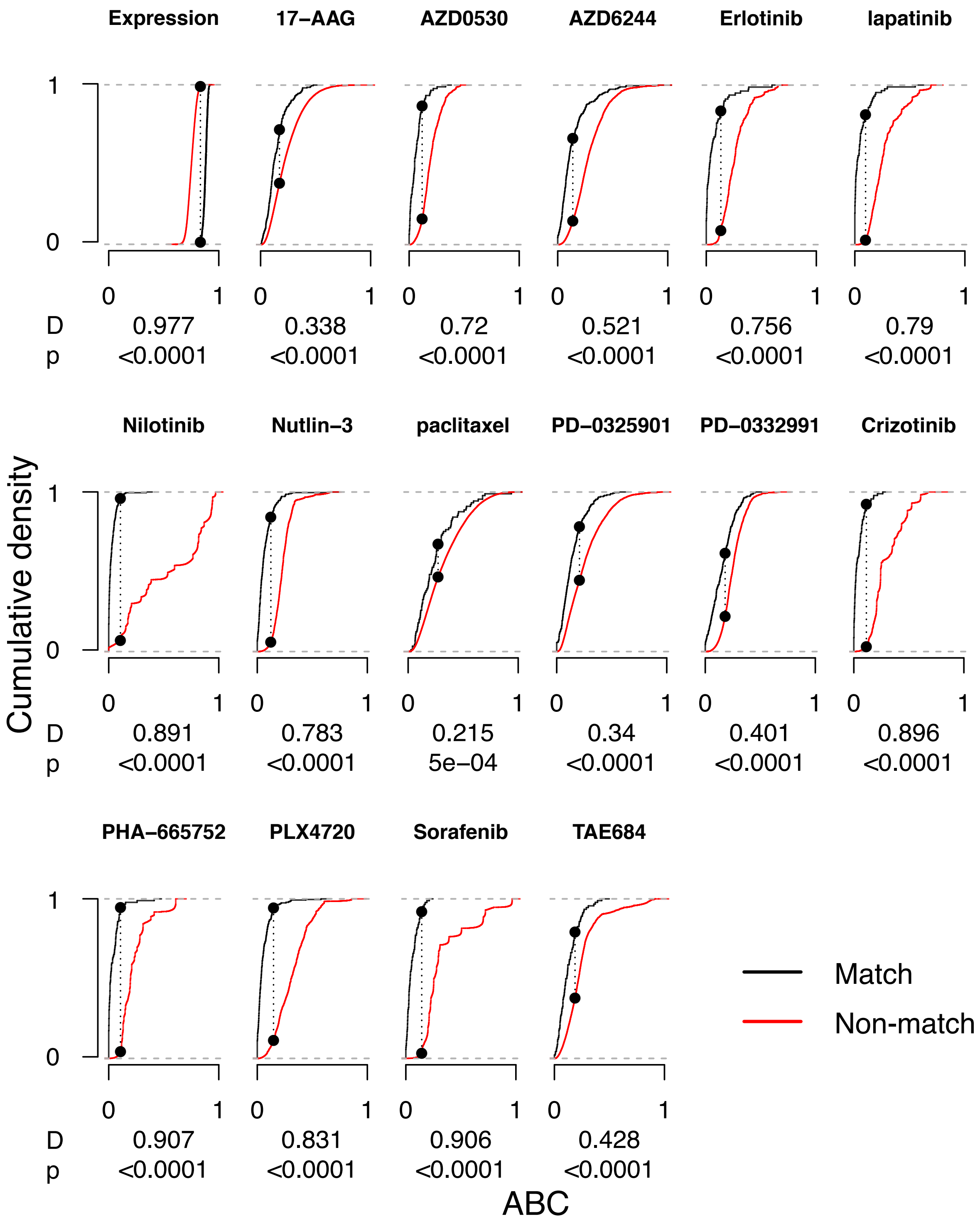


Supplementary Figure 4: ECDF plots of ABC values for all matching and non-matching cell line pairs between GDSC and CCLE per drug. Max distance between the matching and non-matching ECDF curves is represented by D, while the p-value is calculated using the Kolmogorov-Smirnov test.

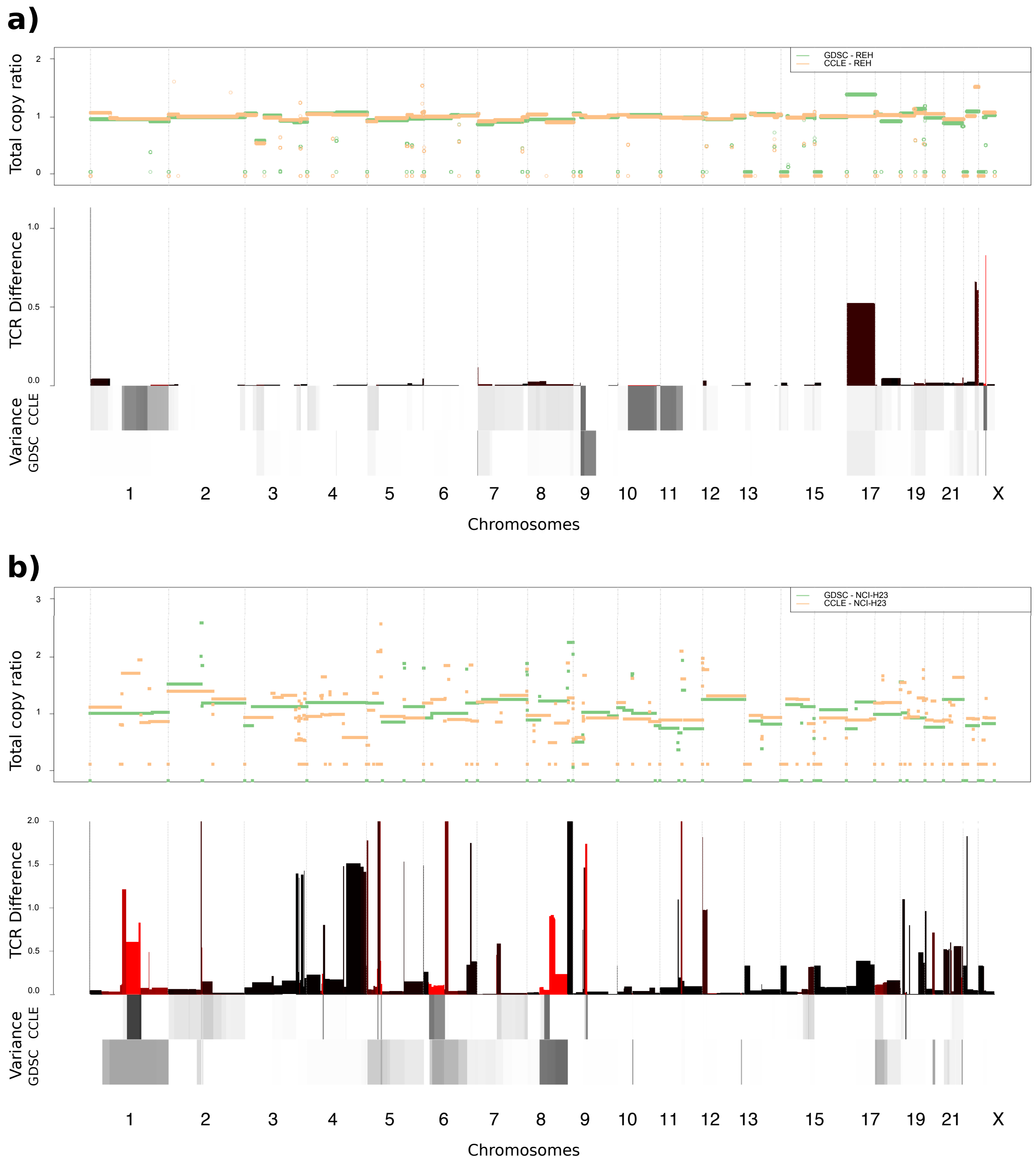


Supplementary Figure 5: Copy number plots for the total copy-ratio between **a)** REH and **b)** NCI-H23. The top track reflects the total CN profile between GDSC (green) and CCLE (orange), while the middle track represents the adjusted total difference between CN profiles where red bars are highly variable CNV-regions. The bottom track is a measure of instability due to pre-processing at a given genomic bin.

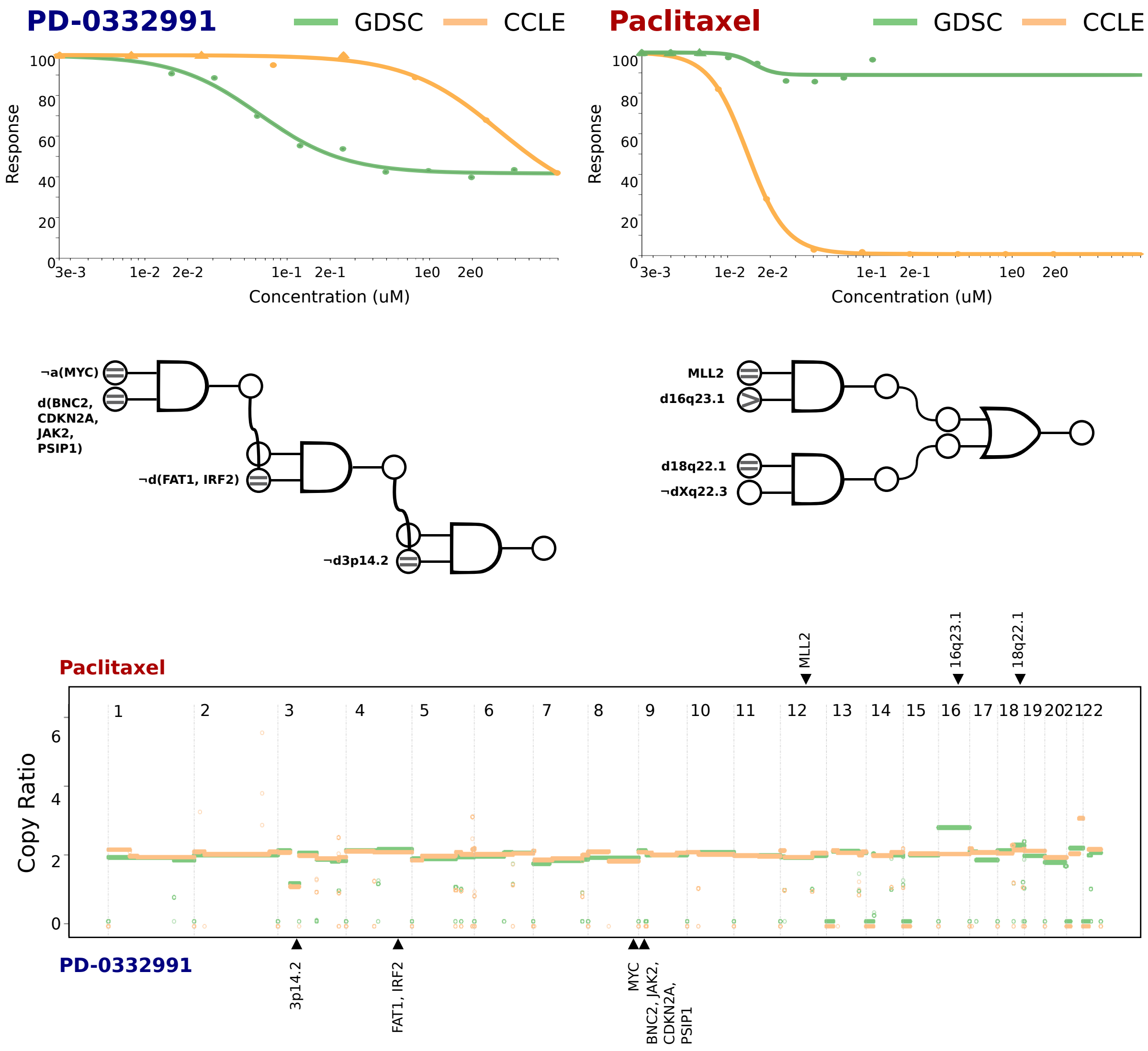


Supplementary Figure 6: Comparison of drug dose-response curves between GDSC and CCLE for the REH cell line. Top panel: Drug dose-response curves between GDSC and CCLE. Middle panel: Logical combinations of top predictive features as estimated by LOBICO, where ">", "=", and "<" would correspond to CN of that feature in GDSC being greater than, equal to, or less than the same feature in CCLE. Bottom panel: The total copy-ratio plot for REH with blue regions indicating predictive features for PD-0332991, and red regions being predictive features for paclitaxel.

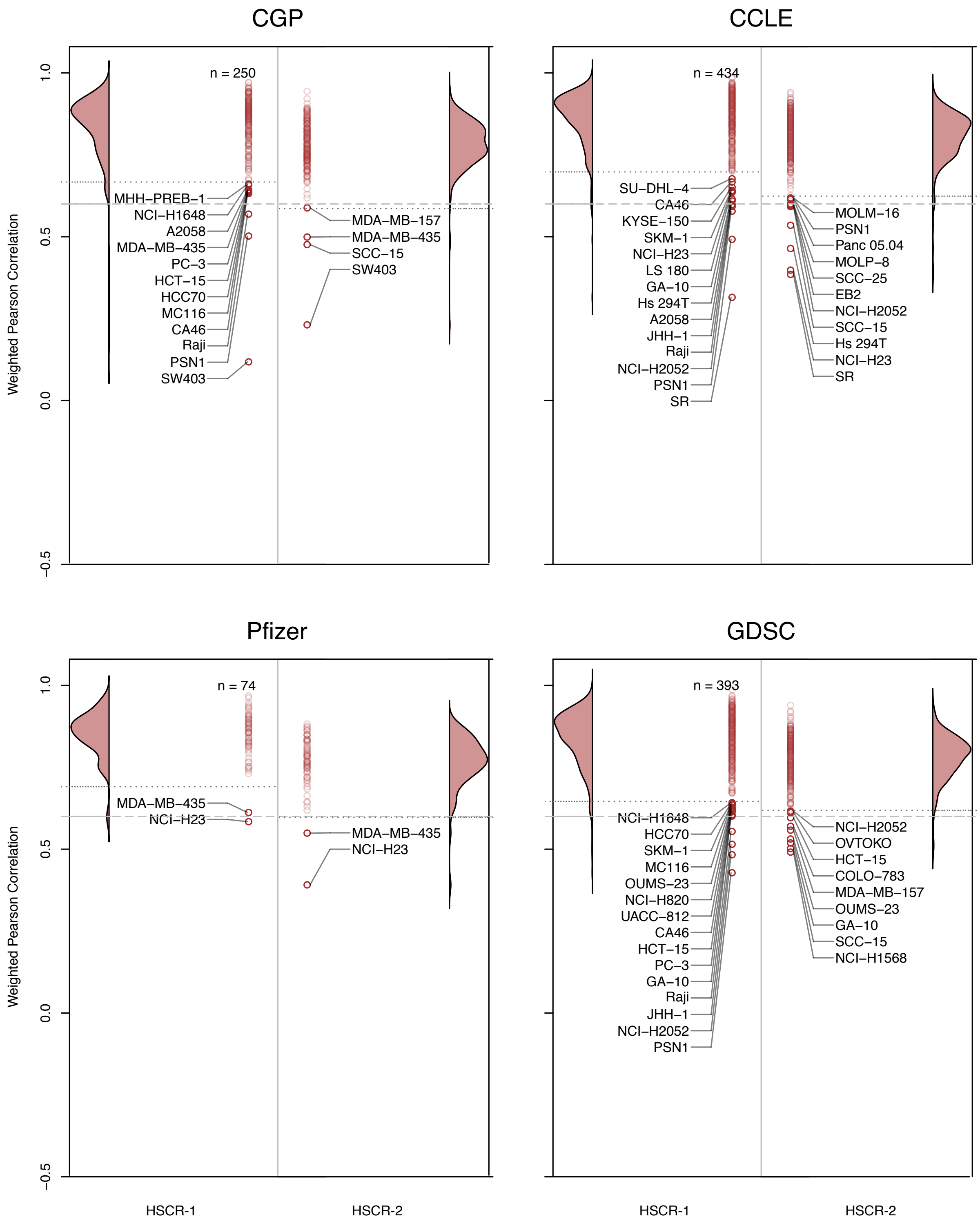


Supplementary Figure 7: Range of Pearson correlation coefficients for HSCRs of all overlapping homonymous cell line pairs between gCSI and CGP, CCLE, Pfizer, and GDSC datasets. Dotted lines indicate the 0.6 hard threshold of discordance, as well as the 3 MADM value. All cell lines below the more stringent threshold are labelled and annotated.

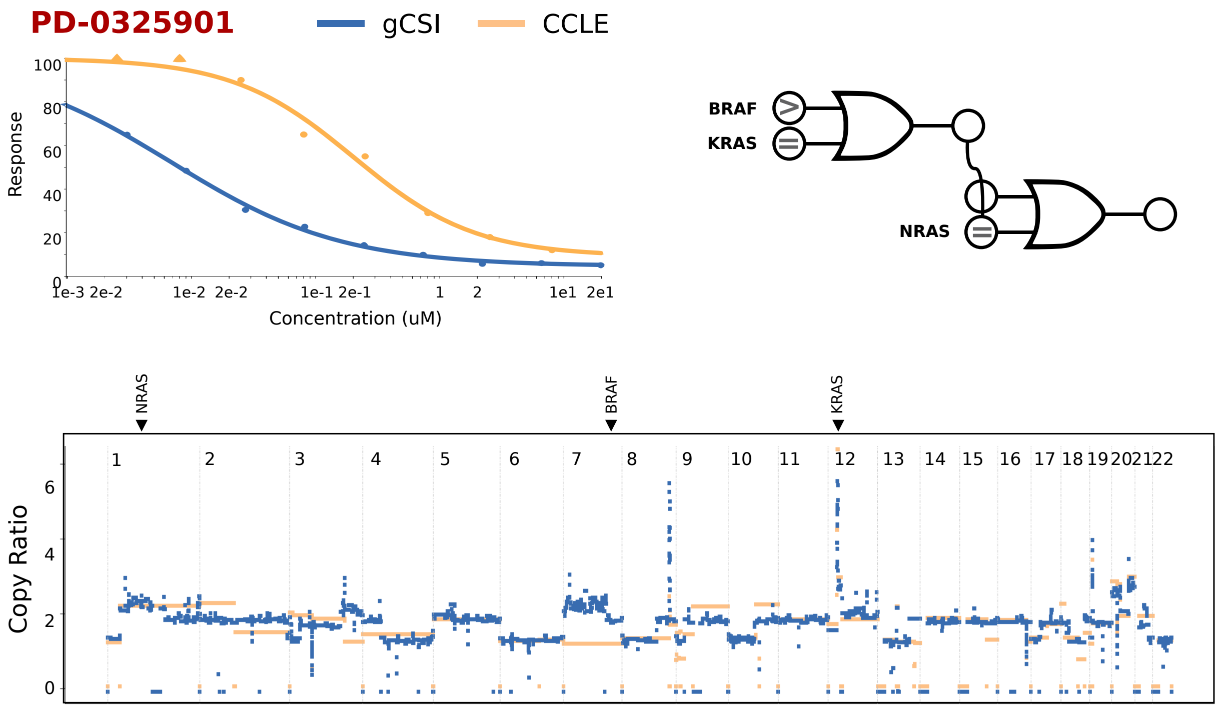


Supplementary Figure 8: Comparison of drug dose-response curves between gCSI and CCLE for the PSN1 cell line. Top-left panel: Drug dose-response curves between gCSI and CCLE. Top-right panel: Logical combinations of top predictive features as estimated by LOBICO, where ">", "=", and "<" would correspond to CN of that feature in gCSI being greater than, equal to, or less than the same feature in CCLE. Bottom panel: The total copy-ratio plot for PSN1 with red regions indicating predictive features for PD-0325901.
