## Supplemental methods for "Karyotypic divergence confounds cellular phenotypes in large pharmacogenomic studies"

### SUPPLEMENTARY METHODS

#### Pharmacogenomic datasets

We analyzed 3,631 Affymetrix Genome-Wide Human SNP 6.0 (Affy6) arrays and 668 HumanOmni2.5-Quad (Omni) arrays from a large compendium of 6 pharmacogenomic studies (Table 1). CEL files and IDAT were downloaded from the accession codes outlined in Table 1 and the cell lines annotations were obtained from PharmacoDB (version 1.0.0) [23,24].

#### Preprocessing of Affymetrix SNP 6.0 arrays

Raw Affy6 files were pre-processed using the affymetrix power tools (APT version 1.16.1) set of tools. As per Affy6 guidelines, samples and datasets were removed based on poorly resolved genotype clusters and a bias towards restriction enzyme digestion during library preparation [25]. The genotypes for all samples in a dataset were called together using the birdseed (version 2) genotyping algorithm implemented in APT with the GenomeWideSNP_6 Annotations Release 35 manifest. Preprocessing of signal intensities were computed in APT through quantile normalization followed by Tukey's median-polish summarization. Homologue-specific copy ratios (HSCR) were then called using HAPSEG (version 1.1.1). Using HSCRs, we then computed the absolute allele-specific somatic copy numbers (ASCN) through the use of ABSOLUTE, a tool that uses a probabilistic method to fit the corresponding HSCRs to a Gaussian mixture model to infer discrete copy numbers [40]. Total copy ratios (TCRs) were called from HAPSEG as the average of both HSCRs at each segment.

#### Preprocessing of Illumina Human Omni Arrays

Raw Omni files were preprocessed using GenomeStudio version 2 using the HumanOmni2.5-Quad version 1.0 manifest and HumanOmni2.5-Quad cluster file. Genotype was called using the Genotype Module. Signal intensities were converted to log-R ratios (LRR) and B-allele frequencies (BAF) using the cnvPartition CNV Analysis Plugin version 3.2.1 which was used as input for ASCAT version 2.5.1 in Tumor-Only mode to generate copy-number profiles.

#### Quality control metrics

Quality of the raw Affy6 data was first assessed using the apt-geno-gc program in affymetrix power tools (version 1.16.1). The contrast quality control (CQC) metric was used to evaluate how well three genotype clusters (AA, AB, BB) could be resolved. A CQC score below 0.4 was used to indicate samples that could not resolve base calls, while a difference between CQC score for Nsp and Sty fragments >2 indicated issues in the efficacy of the restriction enzymes used during sample preparation. Only samples that passed these metrics were kept for further analyses, while datasets that had a mean CQC less than 1.7 were also excluded.

Quality of the raw Omni data was assessed using the QC metrics outlined by the Infinium Genotyping Data Analysis technote. Samples were examined for low quality as defined by having a sample call rate less than 0.90. Additionally, the dataset was manually inspected by scoring visualizing the call rate as a function of sample number, and 10% GenCall (a quality metric calculated for each genotype) as a function of sample call.

#### Comparing genotypes between samples

To compute the concordance between genotypes of two samples using the Affy6 array, we used all 909,622 SNPs to establish concordance between cell lines. To compare genotypes between datasets using the Omni array with those using the Affy6 arrays, we intersected the SNPs based on genomic location and took the concordance of the resulting 337,353 SNPs between all samples. Concordance between the genotype profiles of all pairs of samples, irrespective of the dataset of origin, was estimated using the method outlined by Hong et al. [28].

$$Conc_{i,j} = \frac{1}{N}\sum_{k=1}^{N} n_k, n_k = \begin{cases} 1(G_k^i = G_k^j) \\ 0(G_k^i \neq G_k^j) \end{cases}$$

Where $N$ refers to the population of SNPs and $G_k^i$ is the genotype for SNP $k$ on sample $i$ and $G_k^j$ is the genotype for SNP $k$ on sample $j$.

A single sample was identified to be the same cell line as another or originating from the same individual if they met either of the following criteria: a) they shared >80% genotypic identity, or b) if they were within 3 median absolute deviations away from the median (MADM) for the corresponding “matching cell lines” distribution. These two measures were used as the range of genotype concordance varied depending which datasets were being compared. For instance, many of the comparisons between GDSC and CGP were expected to be of high concordance due to the samples being handled at the same institute or the files being the same, thus a more stringent threshold (MADM) is needed to flag cell lines of interest.

#### Comparing karyotypes between samples

To calculate concordance between copy-number profiles, we resampled without replacement and reprocessed 60% of raw files from each dataset 20 times. Copy number profiles were segmented into 5 kb genomic bins and copy-number profiles and an associated variance-based weight for each bin was estimated. Weights for a genomic bin, $w_b$, were calculated as a function of the standard deviation around the copy-number of the 5 kb genomic bin, $\sigma_b$:

$$w_b = \frac{1}{(\sigma_b + 1)^2}$$

A weighted Pearson correlation coefficient (r) between two copy number profiles, $\rho_{ij}$, was then calculated as:

$$s_i = \frac{\sum_b (w_b^i w_b^j)(x_b^i - m_i)^2}{\sum_b (w_b^i w_b^j)}, \quad s_j = \frac{\sum_b (w_b^i w_b^j)(x_b^j - m_j)^2}{\sum_b (w_b^i w_b^j)}$$

$$s_{ij} = \frac{\sum_b (w_b^i w_b^j)(x_b^i - m_i)(x_b^j - m_j)}{\sum_b (w_b^i w_b^j)}$$

$$\rho_{ij} = \frac{s_{ij}}{\sqrt{s_i s_j}}$$

Where $b$ is each 5 kb genomic bin, $w_b^i$ is the weight for bin $b$ in sample $i$, and $x_b^i$ is the copy-number ratio for bin $b$ in sample $i$. Similar to genotype concordance scores, samples were identified to have the same karyotype if they had a >60% correlation or were within 3 MADMs of the corresponding “matching cell lines” distribution.

#### Cellular phenotypes

Reprocessed drug curves and gene expression data were accessed from the R PharmacoGx (version 1.10.3) package [23]. Expression space was represented by the landmark genes defined in the L1000 dataset [29]. The recomputed area above the curve (AAC) were used as measurements of drug sensitivity, with higher AACs corresponding to higher sensitivity [22]. Distances between drug curves were computed as the area between the curve (ABC) fitted on overlapping drug concentrations between any two datasets (Figure 1).

#### Comparing phenotypes

To build a reference distribution of what is an expected amount of variation drug sensitivity for any given cell line, we built a "Matching" distribution consisting of the delta AAC for all cell lines pairs with matching annotations for drug $d$. Similarly, the expected amount of difference in drug sensitivity measurements was estimated using a "Non-matching" distribution which consisted of the delta AAC for all cell lines with non-matching annotations for drug $d$. To adjust for the many resistant cell lines for a given drug skewing our non-matching distribution, we removed all cell line pairs in the non-matching that were both resistant given that:

$$r_i \wedge r_j < 0.2 \mid d \neq \text{paclitaxel}$$

$$r_i \wedge r_j < 0.4 \mid d = \text{paclitaxel}$$

In order to determine whether a given cell line pair had a comparable phenotype, we first used a Kolmogorov-Smirnov test to ensure that the matching and non-matching distributions were statistically different. By using a kernel density estimation to approximate the probability density function or both distributions, we then calculated the log likelihood ratio of non-matching to matching for the cell line pair of interest
